# Coupling machine learning and epidemiological modelling to characterise optimal fungicide doses when fungicide resistance is partial or quantitative

**DOI:** 10.1101/2022.09.17.508365

**Authors:** Nick P Taylor, Nik J Cunniffe

**Author notes:** **For correspondence** (NPT).

## Abstract

Increasing fungicide dose tends to lead to better short-term control of plant diseases. However, high doses select more rapidly for fungicide resistant strains, reducing long-term disease control. When resistance is qualitative and complete – i.e. resistant strains are unaffected by the chemical and resistance requires only a single genetic change – using the lowest possible dose ensuring sufficient control is well-known as the optimal resistance management strategy. However, partial resistance (where resistant strains are still partially suppressed by the fungicide) and quantitative resistance (where a range of resistant strains are present) remain ill-understood. Here we use a model of quantitative fungicide resistance (parameterised for the economically-important fungal pathogen *Zymoseptoria tritici*) which handles qualitative partial resistance as a special case. We show that – for both qualitative partial resistance and quantitative resistance – although low doses are optimal for resistance management, for some model parameterisations the benefit does not outweigh the improvement in control from increasing doses. Via a machine learning approach (a gradient-boosted trees model combined with Shapley values to facilitate interpretability) we interpret the effect of parameters controlling pathogen mutation and characterising the fungicide, in addition to the timescale of interest.

## Introduction

Plant pathogens have a significant impact on global food production (***Strange and Scott, 2005***; ***Ristaino et al., 2021***). Diseases caused by plant pathogens routinely lead to large losses in crop yields (***Savary et al., 2019***); an estimated 20% of losses of global crop production are caused by disease (***Jorgensen et al., 2017***). However, more than 900 million people are undernourished (***Rockström et al., 2020***), and food production will need to increase by an estimated 60% by 2050 (***Ristaino et al., 2021***). Fungal plant diseases can be particularly damaging due to the potential for prolific spore production, long-range spore dispersal, high mutation rates and potential for both sexual (***Möller and Stukenbrock, 2017***) and asexual reproduction (***Fones et al., 2020***). Further, fungal pathogens are strongly influenced by climatic factors, so changes in climate are likely to affect the incidence and severity of many diseases caused by fungi (***Lucas, 2017***). Modern agricultural ecosystems offer reduced species diversity when compared to natural ecosystems, but higher host and pathogen density, creating highly conducive environments for rapid dispersal and evolution of fungal plant pathogens (***McDonald and Stukenbrock, 2016***). Current control mechanisms rely strongly on chemical control from fungicides. However, their control is regularly challenged by fungicide resistance due to the enormous evolutionary potential of fungal pathogens.

Control of fungal pathogens is extremely economically important; approximately 16 billion US dollars are spent every year on fungicides globally (***Jorgensen et al., 2017***). Mathematical modelling can be an invaluable tool enabling us to understand the complicated mechanisms underlying fungicide resistance development (***Corkley et al., 2022***; ***Cunniffe et al., 2015***). Theoretical and modelling studies offer numerous advantages over experimental studies. For example, field trials are frequently very expensive and it can be difficult to control for confounding factors such as environmental variability between years. The time-scales involved in fungicide resistance studies can be extremely long, which leads to increased cost and enormous delays in obtaining results when compared to theoretical studies. Further, fungicide resistance is often (at least initially) present at frequencies so low where it can be difficult to obtain accurate measurements.

Most fungicide resistance modelling studies address qualitative resistance (i.e. where a single mutation causes fungicide resistance) (***Hobbelen et al., 2011a***,b, ***2013***; ***Mikaberidze et al., 2017***; ***Elderfield et al., 2018***; ***Mikaberidze et al., 2014***; ***van den Berg et al., 2013***; ***Taylor and Cunniffe, 2022b***). However, many contemporary fungicides in use (e.g. the azole and succinate dehydrogenase inhibitor (SDHI) fungicides (***Kirikyali et al., 2017***; ***Torriani et al., 2015***)) are challenged by quantitative resistance (i.e. where several successive mutations are needed to acquire considerable levels of resistance, meaning many strains can be present each with their own level of resistance (***Mikaberidze et al., 2017***; ***McGrath, 2007***; ***Didelot et al., 2016***)). Many questions in the fungicide modelling literature about resistance management have only been addressed in the case of qualitative resistance.

One of the key questions about fungicide resistance management concerns fungicide dose; when should low doses be used in favour of high doses? The vast majority of experimental and modelling literature suggests that high doses select more strongly for qualitative resistance (***van den Bosch et al., 2011***), due to increased selection pressure in line with the so-called governing principles of fungicide resistance (***van den Bosch et al., 2014***). This means that low doses are often preferable when tackling qualitative resistance, so long as adequate yield can be maintained.

In theory high doses may be useful in targeting diploid pathogens (***van den Bosch et al., 2011***). To illustrate this, consider a population containing homozygote sensitive (denoted SS), homozygote resistant (denoted RR) and heterozygote individuals (denoted RS). If resistance is rare, and mating random, then it is most likely that the SR and RR strains will mate with a homozygote sensitive (SS) individual, resulting in more homozygote sensitive or heterozygote individuals. The heterozygote individuals can be better controlled by high doses which can remove resistant alleles from the population (***van den Bosch et al., 2011***). However, most fungal pathogens are haploid or largely clonal meaning that low doses are usually recommended (***van den Bosch et al., 2014***, ***2011***).

Partial resistance occurs when the resistant strain is still partially suppressed by the fungicide (as opposed to complete resistance, where the resistant strain is completely unaffected by the fungicide). If dose-response curves converge at high doses, which might occur with partial resistance, then high doses could result in reduced selection for resistance (***van den Bosch et al., 2014***, ***2011***; ***Neve and Powles, 2005***). Dose-response convergence means that the difference between growth rates of resistant and sensitive strains is smaller for high doses than at smaller or intermediate doses. Whether this happens depends on the shape of the dose-response curve. This can lead to reduced selection for fungicide resistance at high doses compared to lower doses.

Models often address the so-called ‘emergence phase’ or the ‘selection phase’ of fungicide resistance. The emergence phase is when resistance arises in a previously sensitive population through mutations (***van den Bosch and Gilligan, 2008***). The selection phase is when resistant genotypes are present in a population and subsequently selected for by fungicide use (***van den Bosch and Gilligan, 2008***). Most modelling studies focus solely on either the emergence phase (***Mikaberidze et al., 2017***) or the selection phase (***Hobbelen et al., 2011a***,b, ***2013***). Two types of partial qualitative resistance are described in ***Mikaberidze et al***. (***2017***): Type 1 where resistance is characterised in terms of the maximum fungicide effect; Type 2 where resistance is characterised in terms of the ‘slope’ of the fungicide effect with dose. For Type 1 partial resistance, high doses were found to accelerate resistance emergence, whereas for partial resistance Type 2 there was a minimum in emergence time for intermediate doses, suggesting low or high doses should be favoured. The driver of the result is whether the dose-response curves converge at high doses, which happens with partial resistance Type 2 but not with Type 1.

Although ***Mikaberidze et al***. (***2017***) address partial resistance, the model targets only qualitative resistance, and so considers two pathogen strains only. This means that the model does not adequately address the wide range of pathogen strains possible in the quantitative resistance case. However, dose-convergence is possible when resistance is quantitative as well as when it is qualitative but partial, meaning that high doses may result in reduced selection at higher doses. While ***Shaw*** (***1989***) defined a theoretical model of polygenically controlled (i.e. quantitative) fungicide resistance, it was not fitted to field data and did not have a notion of crop yield. The model in ***Taylor and Cunniffe*** (***2022a***) addresses quantitative resistance parameterised for control of Septoria tritici blotch (caused by *Zymoseptoria tritici*), the most prevalent disease of wheat worldwide (***Suffert et al., 2011***), using the azole fungicide ‘prothioconazole’. In ***Taylor and Cunniffe*** (***2022a***) applications were always at full dose; in this work we extend the model to consider lower doses and explore the effect of dose choice on disease control and resistance management. We also consider a range of epidemiological and fungicide parameter choices, to allow us to explore beyond the wheat-septoria/azole system.

Many models of fungicide resistance development neglect to explicitly model pathogen mutation (***Hobbelen et al., 2011a***,b, ***2013***; ***Mikaberidze et al., 2017***; ***Elderfield et al., 2018***; ***Mikaberidze et al., 2014***), despite its important evolutionary role (***McDonald et al., 2022***). The model in ***Taylor and Cunniffe*** (***2022a***) includes pathogen mutation and is capable of modelling both emergence of resistance and selection for resistant strains present in the population. We will explore the effect of parameters controlling the pathogen mutation rate and mutation scale on the optimal dose recommendation.

Some previous plant disease modelling work makes use of sensitivity analyses to explore the effects of different model parameters on the model output. In ***Rimbaud et al***. (***2018***), polynomial regression was used to characterise the epidemiological model output in terms of 6 model parameters. Sobol’s method was used to evaluate which features have the strongest impact on the model. In this work we introduce an alternative method, common in machine learning, to explain the effect of 8 model parameters on the output of the quantitative resistance model. Machine learning approaches are becoming increasingly important in a wide variety of applications ranging from fraud detection to speech recognition. Gradient-boosted trees models are one such algorithm capable of excellent results on a wide range of problems, often outperforming polynomial regression models, random forests and neural networks (***Chen and Guestrin, 2016***). We combine a gradient-boosted trees model with Shapley values (***Shapley, 1953***; ***Lundberg et al., 2020***), a technique from game theory for allocating importance to members of a coalition (in our case, variables in a model). This allows us to rank and understand the impact of different parameters on the output of our model, i.e. the optimal dose. The approach facilitates model interpretability and explanation whilst making use of the accuracy of gradient tree boosting.

In this paper we address the following questions:

- Can high fungicide doses ever outperform low doses?
- How does this depend on whether fungicide resistance is qualitative or quantitative?
- What is the effect of fungicide efficacy and pathogen mutation on optimal dose choice over different timescales?
- Do Shapley values help make a complex parameter scan (of the type common in plant disease epidemiology) more interpretable?
- What mechanisms cause optimality of low (or high) doses?

## Methods

### Model explanation

We use the model of quantitative fungicide resistance from ***Taylor and Cunniffe*** (***2022a***). It addresses a diverse pathogen population containing strains which have different sensitivities to a fungicide. These are described by a continuous ‘trait value’ *k* taking values in [0, 1], where *k* = 0 corresponds to a fully sensitive pathogen strain (completely suppressed by a fungicide application) and *k* = 1 corresponds to a fully resistant pathogen strain (completely unaffected by a fungicide application).

The model tracks healthy/susceptible leaf tissue (denoted by *S*(*t*), where *t* is the time within a single growing season) and leaf tissue infected by strain *k* (denoted by *I* (*k*, *t*)). The model is applied over multiple growing seasons. It includes mutation, a host growth function *g*(*t*) and a senescence function Γ(*t*). The infection rate *β*(*k*, *t*) depends on the pathogen strain *k*, since each strain responds differently to a fungicide application. We refer the reader to ***Taylor and Cunniffe*** (***2022a***) for a full explanation of the model derivation and details on the model fitting process. Although the full form of the fitted model as presented in ***Taylor and Cunniffe*** (***2022a***) included the effect of disease-resistant crop varieties, for simplicity here we do not model host plant protection, meaning we use the so-called ‘fungicide only’ form of the model from ***Taylor and Cunniffe*** (***2022a***).

The model is defined as:

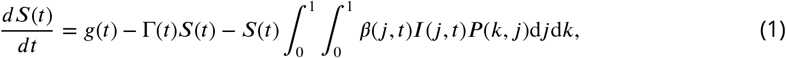

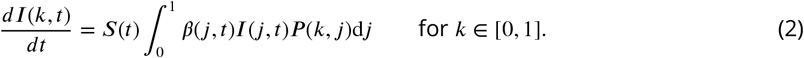

Variable and function definitions are given in Table 1, and the functions *g*(*t*), Γ(*t*), *P*(*k*, *j*) and *β*(*k*, *t*) are described in more detail below. Parameter values as fitted in ***Taylor and Cunniffe*** (***2022a***) are found in Appendix 1 Table 1.

**Table 1.**
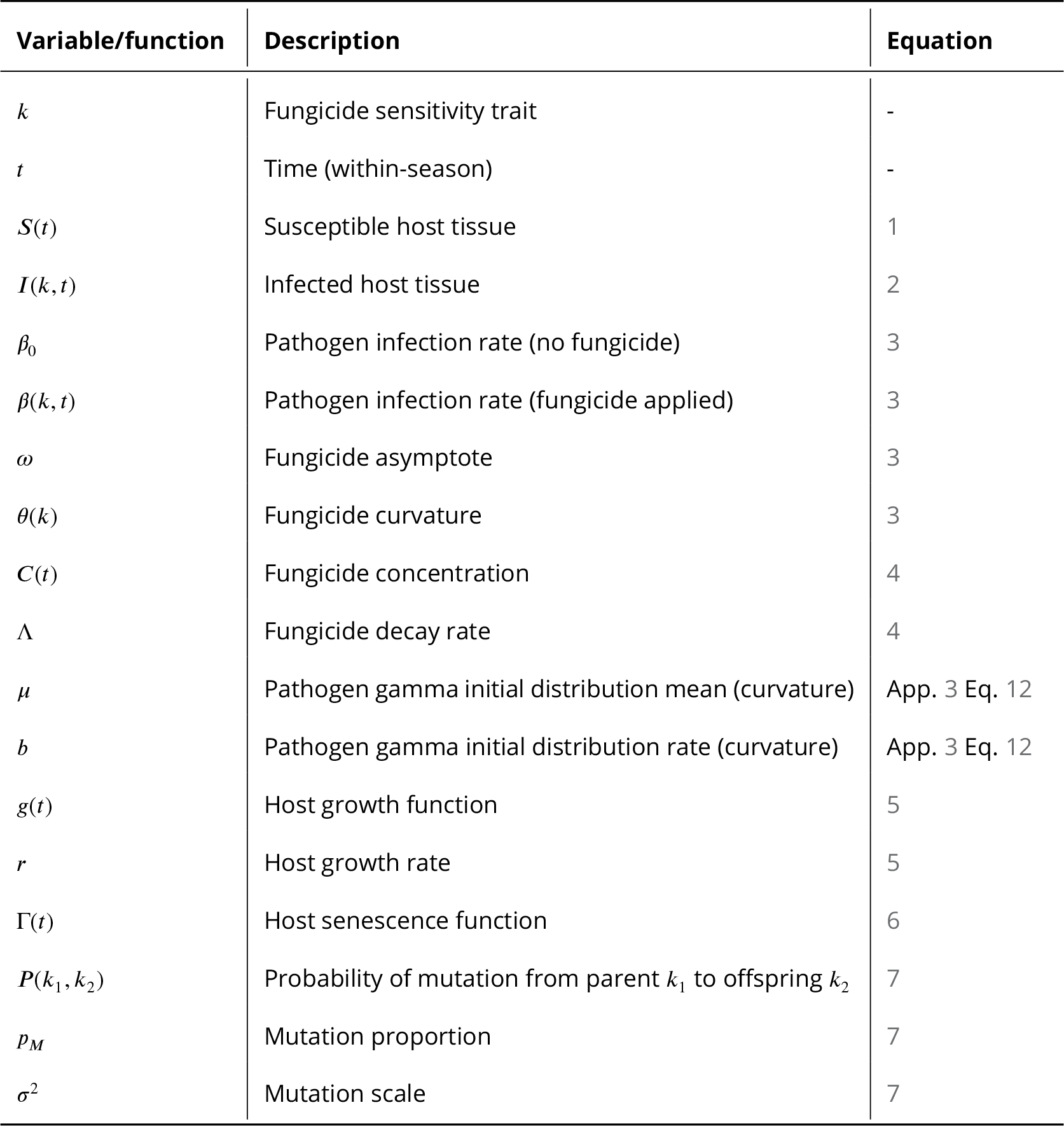
List of variables and functions used in the model. Default values are given in Appendix 1 Table 1.

| Variable/function | Description | Equation |
| --- | --- | --- |
| $k$ | Fungicide sensitivity trait | - |
| $t$ | Time (within-season) | - |
| $S(t)$ | Susceptible host tissue | 1 |
| $I(k, t)$ | Infected host tissue | 2 |
| $\beta_0$ | Pathogen infection rate (no fungicide) | 3 |
| $\beta(k, t)$ | Pathogen infection rate (fungicide applied) | 3 |
| $\omega$ | Fungicide asymptote | 3 |
| $\theta(k)$ | Fungicide curvature | 3 |
| $C(t)$ | Fungicide concentration | 4 |
| $\Lambda$ | Fungicide decay rate | 4 |
| $\mu$ | Pathogen gamma initial distribution mean (curvature) | App. 3 Eq. 12 |
| $b$ | Pathogen gamma initial distribution rate (curvature) | App. 3 Eq. 12 |
| $g(t)$ | Host growth function | 5 |
| $r$ | Host growth rate | 5 |
| $\Gamma(t)$ | Host senescence function | 6 |
| $P(k_1, k_2)$ | Probability of mutation from parent $k_1$ to offspring $k_2$ | 7 |
| $p_M$ | Mutation proportion | 7 |
| $\sigma^2$ | Mutation scale | 7 |

#### Infection rate / effect of fungicide

Denote the fungicide concentration at time *t* by *C*(*t*). Then the infection rate, *β*(*k*, *t*), of strain *k* at time *t* as defined in ***Taylor and Cunniffe*** (***2022a***) is:

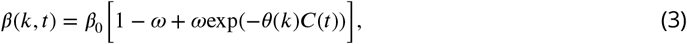

where *θ*(*k*) = −log(*k*) is the so-called fungicide ‘curvature’ parameter for strain *k*, and *ω* is the fungicide asymptote parameter. ***Taylor and Cunniffe*** (***2022a***) only considered the case where *ω* = 1. When the concentration is 0, the exponential term is 1, so when there is no fungicide present all strains behave identically. When the fungicide is applied the infection rates of strains with lower *k* values are suppressed more than that of strains with higher *k* values.

The fungicide concentration *C*(*t*) is assumed to decay exponentially after each application with time *t* at rate Λ (Appendix 1 Table 1). The fungicide concentration equation is:

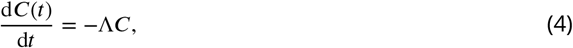

where *C* is initially 0 but increases instantaneously by an amount *D* every time a dose of *D* of the fungicide is applied. In ***Taylor and Cunniffe*** (***2022a***), only a full dose of 1 was considered. In this work we consider lower doses, using 10 dose choices: 0.1,0.2, … 1. In this work we restrict our attention to fungicide application programmes containing 2 sprays at the spray times conventionally called *T*_2_ and *T*_3_ (Table 2) (***van den Berg et al., 2013***).

**Table 2.**
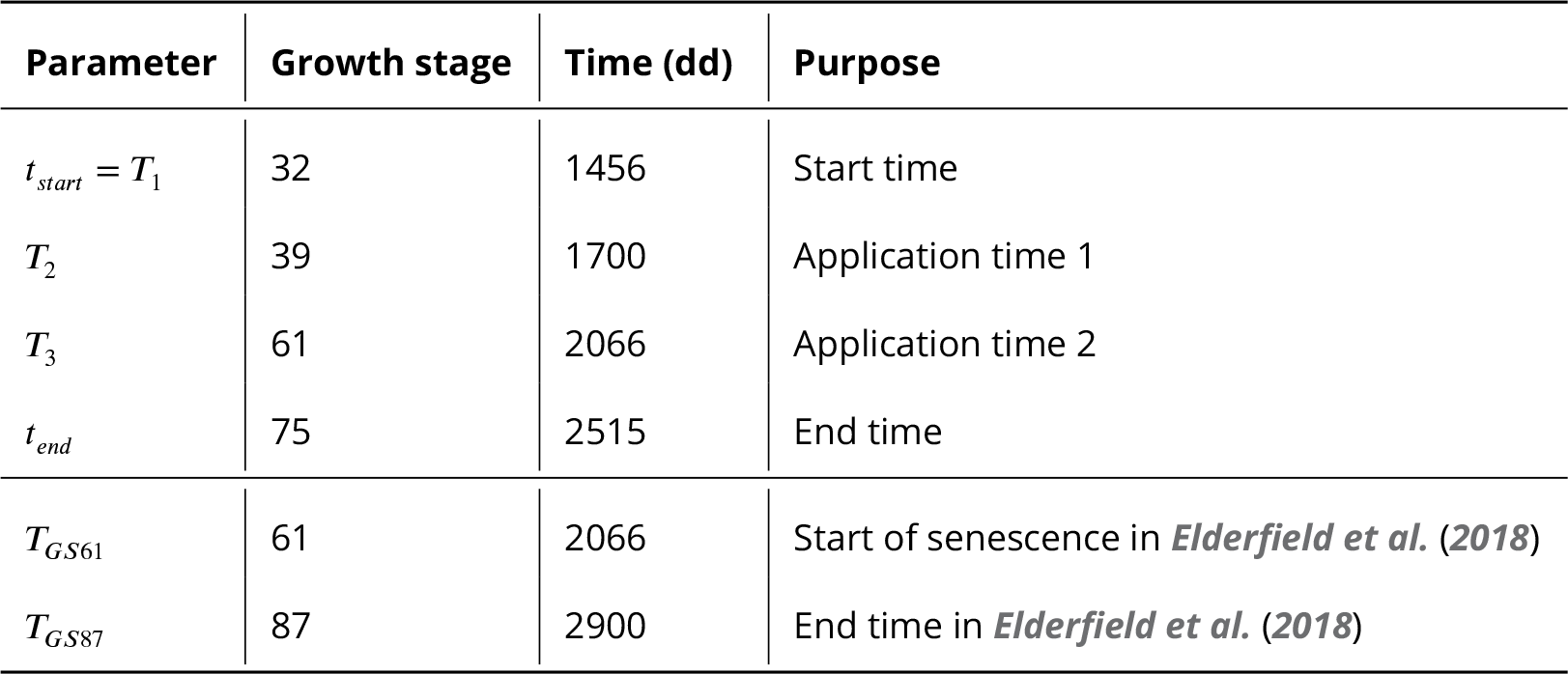
Model running times, as found in ***Taylor and Cunniffe*** (***2022a***). Note that ‘dd’ refers to the units ‘degree-days’. Degree-days are calculated based on a base temperature of 0°C, and the growing season average temperature in Cambridge (UK) during 1984 to 2003 of 15.2°C (i.e. one calendar day equals 15.2 degree-days), as in ***van den Berg et al***. (***2013***). Note that the start time is at growth stage 32, and the host growth and epidemic progress until this point is scaled into the model initial conditions as in ***Taylor and Cunniffe*** (***2022a***).

| Parameter | Growth stage | Time (dd) | Purpose |
| --- | --- | --- | --- |
| $t_{start} = T_1$ | 32 | 1456 | Start time |
| $T_2$ | 39 | 1700 | Application time 1 |
| $T_3$ | 61 | 2066 | Application time 2 |
| $t_{end}$ | 75 | 2515 | End time |
| $T_{GS61}$ | 61 | 2066 | Start of senescence in *Elderfield et al. (2018)* |
| $T_{GS87}$ | 87 | 2900 | End time in *Elderfield et al. (2018)* |

#### Host growth and senescence

Within the season, the host is assumed to initially grow before a period of senescence. We use the same forms for growth, and for senescence, as in ***Hobbelen et al***. (***2011b***); ***Elderfield et al***. (***2018***); ***Taylor and Cunniffe*** (***2022b***). Let *A*(*t*) be the total amount of tissue (healthy and infected). Then the growth equation is:

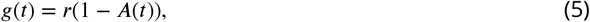

where *r* is the host growth rate (constant, see Appendix 1 Table 1).

The (time-dependent) rate at which leaf senescence occurs, is:

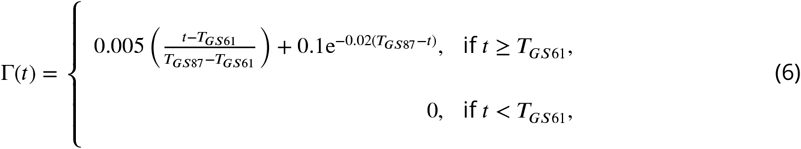

as in ***Hobbelen et al***. (***2011a***); ***Elderfield et al***. (***2018***); ***Taylor and Cunniffe*** (***2022b***,a) (Table 2).

#### Mutation

We incorporate pathogen mutation, meaning that a small proportion of each pathogen strain’s offspring takes a different trait value. ***Taylor and Cunniffe*** (***2022a***) used a Gaussian mutation kernel, assuming mutation events occur with probability *p*_*M*_ and with mutation scale *σ*^2^. Let *δ*(*x*) be the Kronecker delta (which is 1 when *x* = 0 and 0 otherwise). Then for a strain with trait value *j*, we denote the probability *P*(*k*, *j*) of its offspring taking trait value *k*:

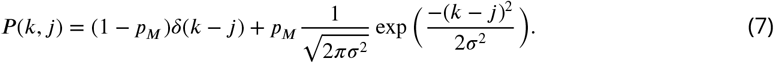

Any offspring predicted to take negative trait values are instead given trait value 0 and any that are predicted to take values greater than 1 are given trait value 1.

Offspring with trait value *k* has a per capita growth rate that is found by integrating over all possible parents with trait value *j*:

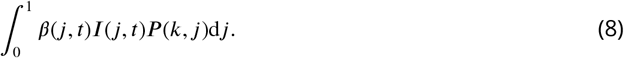

In the equation for 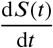, (Equation 1), we integrate again to capture the per capita growth of all strains in the population.

#### Initial distribution of trait values

To describe the initial distribution of trait values, we use a gamma distribution for the curvature values (equivalent to −log(*k*)), as in ***Taylor and Cunniffe*** (***2022a***). After each growing season, the population is normalised so that the total inoculum when integrating over all strains is *I*_0_ in the next season (Appendix 1 Table 1). However, the relative amounts of each strain changes due to the fungicide applications.

### Qualitative resistance

The quantitative/polygenic resistance model we present can entirely capture the behaviour of a qualitative/monogenic system simply by choosing an initial pathogen distribution which has density at two *k* values only and setting the mutation proportion *p*_*M*_ to be 0.

Denote these two pathogen strains *k*_*r*_, *k*_*s*_, at proportions *p*_*s*_, 1 − *p*_*s*_. Then the mean trait value is:

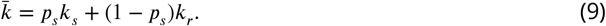

Using the model in this way allows us to directly compare the optimal dose choice for a qualitative system to a quantitative system, using the same model. We complete have flexibility over the choices of *k*_*r*_ and *k*_*s*_, which allows us to explore a range of different fungicide efficacies against resistant strains and different levels of partial resistance. Initially we will explore the behaviour of the qualitative/monogenic system with partial resistance before comparing to the results of the quantitative system.

### Parameter scan

In order to analyse when low doses perform better than high doses, we ran a parameter scan across 10,000 model runs. Values were sampled randomly and independently for a variety of parameters, allowing us to explore the behaviour of the model for a range of parameterisations. For each run in the ensemble, we sampled values for: the two fungicide initial distribution parameters; the fungicide dose response parameters (decay rate and asymptote); and the two mutation parameters (scale and proportion) – see Table 3. We also calculate the initial mean effect of the fungicide at full dose, *ν*, which is the ‘maximum effect’ of the fungicide at the start of the first season. This depends on the fungicide distribution mean 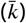 and asymptote (*ω*) as follows:

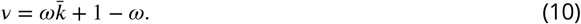

**Table 3.** Parameters/features involved in the parameter scan. The first six are randomly and independently sampled for each parameter run, while the mean fungicide effect at full dose (*ν*) is calculated based on these, and year varies between 1 and 35 in each run. The first four parameters are sampled from a uniform distribution, and the two mutation multipliers are sampled from a log-uniform distribution. The multipliers for decay rate, mutation scale and mutation proportion relate to the default values for Λ, *σ*^2^ and *p*_*M*_ (Appendix 1).

| Symbol | Description | Range | Equation |
| --- | --- | --- | --- |
| $\mu$ | Pathogen gamma initial distribution mean (curvature) | [0, 25] | App. 3 Eq. 12 |
| $b$ | Pathogen gamma initial distribution rate (curvature) | [0, 5] | App. 3 Eq. 12 |
| $\omega$ | Fungicide asymptote | [0, 1] | 3 |
| $M_d$ | Fungicide decay rate multiplier | $\left[\frac{1}{3}, 3\right]$ | 4 |
| $M_s$ | Mutation scale multiplier | [0.1, 10] | 7 |
| $M_p$ | Mutation proportion multiplier | [0.1, 10] | 7 |
| $\nu$ | Mean fungicide effect at full dose | Depends on $\omega$, $\mu$, $b$ | 10, 11 |
| $Y$ | Year in model run | [1, 35] (but not sampled) | - |

This means that when *ω* = 1, 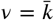, and when *ω* = 0, *ν* = 1. Note that 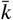 can be written in terms of *μ*, *b* (see Appendix 3):

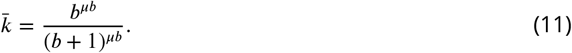

For each run in the ensemble, we test out 10 doses from 0.1 to 1, allowing each dose strategy to be used for 35 years. The model returns the yield in each year, and in a given year *N* we are interested in the ‘best dose’, i.e. the dose which gives the highest yield in year *N* given the use of that dose in years 1, 2, … , *N*. The result of the scan was a dataframe containing one row per year of each ensemble member, along with the various variables described above and in Table 3, and the best dose.

### Gradient-boosted trees model

We fitted a gradient-boosted trees model in order to analyse the results of the parameter scan and seek relationships between the different features (i.e. the parameters in Table 3) and the best dose. Gradient-boosted trees models are a machine learning technique based on an ensemble of decision trees. Each decision tree splits the input space into various disjoint regions, and the decision tree output is constant in any of these regions. The ensemble of decision trees is built up so that each successive model is fitted to the residuals of the previous ensemble model. When the loss function is the squared error, the residuals are proportional the derivative (gradient) of the loss function, hence the name gradient-boosting. The ‘learning rate’ (*LR*) is used to limit the impact of any individual tree within the ensemble by scaling the output by a value *LR* ∈ [0, 1]. This leads to much improved accuracy and better generalisation to unseen data. We used the freely-available python package XGBoost (***Chen and Guestrin, 2016***) to fit the gradient boosted regression model to the data. Gradient-boosted trees models give outstanding results across a wide range of applications; 17 of the 29 of the challenge winning solutions to Kaggle problems in 2015 used XGBoost (***Chen and Guestrin, 2016***). They can be more accurate than neural networks and more interpretable than linear models (***Lundberg et al., 2020***). Further, they are capable of dealing with strongly interacting and/or co-linear features.

#### Model fitting

To fit the model we split our data into two parts; a training and test set. These were split in an 80:20 proportion so that the first 8000 runs were used for model training. We optimised for the following hyperparameters (Appendix 2 Tables 1, 2): max depth, number of estimators, learning rate, sub-sample, column sample by tree. This optimisation was performed on the training set, using a 5-fold cross-validation to assess the performance of each set of hyperparameters. The best model was then checked on the test set to check that the model performance did not degrade significantly on completely unseen data. Choosing appropriate hyperparameter values helps avoid overfitting the model but ensures we retain sufficient complexity to capture the relationships found within the data. Model performance (in terms of root mean squared residuals) is shown in Appendix 2 Table 3 and on predictions on (previously unseen) test data in Appendix 2 Figures 1, 2.

### Shapley values

Shapley additive explanations (SHAP, (***Lundberg et al., 2020***)) allow us to explain the output of our gradient-boosted trees model. Shapley values (***Shapley, 1953***) give an indication of how important each input feature is in controlling the value of a given output. They are a method from cooperative game theory which allow us to allocate credit to individual ‘players’ (i.e. model features in our case) for the output of the model (i.e. best dose in our case). The ‘game’ is the task of predicting a single instance (i.e. a single row) of the dataset given feature values. The ‘gain’ is the actual prediction for this instance minus the average prediction for all instances. A ‘coalition’ is a subset of players/features. The Shapley value of a particular feature is a way of measuring that feature’s contribution to the model output, i.e. that player’s contribution to the outcome of the game. For a given feature value and coalition excluding that feature, we can find the marginal contribution of that feature to the prediction when it is added to the coalition, by comparing the prediction from the coalition size *k* averaged over all possible values of the *N* − *k* exluded variables to the prediction from the coalition size *k* + 1 that includes the feature. Then the Shapley value is the (weighted) average over all marginal contributions of that feature value, i.e. the average (over all coalitions excluding the feature) of the change in prediction that occurs when that feature value is included in the coalition. They are a valuable tool facilitating interpretability and explainability of complex and otherwise opaque models.

In general, Shapley values can only be approximated since computing them exactly is NP-hard. However, for gradient-boosted trees, they can be calculated exactly, and in polynomical time (***Lundberg et al., 2020***). We use the open-source Python package ‘shap’ to do so. This helps us analyse the gradient-boosted trees model output and further understand the relationship between the various fungicide parameters and the corresponding best dose across the full range of timescales considered (1-35 years).

## Results

### Can high doses ever be best with qualtitative partial resistance?

If resistance is complete (i.e. the *k* value of the resistant strain is *k*_*r*_ = 1), lower doses are preferable in terms of yield/control to high doses (Figure 1A-D), apart from in the very early years when the resistant proportion is at a very low density. This is because higher doses select more strongly for the resistant strain (Figure 1C), and the fungicide is completely unable to control the resistant strain, leading to reduced yields from higher doses once resistance has developed (Figure 1D). However, if resistance is partial, there are cases where high doses are better over certain time frames (Figure 1E-H). In the very short and very long term, the pathogen population is almost entirely sensitive and (partially) resistant respectively. In both these cases, higher doses offer better control. The trade-off occurs in the intermediate period where resistance develops more quickly (Figure 1G) to the higher dose fungicide programmes, leading to lower yield for higher doses in this intermediate period of time (Figure 1H). When the partial resistance is weak (Figure 1I), it is possible for high doses to always outperform low doses (Figure 1I-L). Although higher doses still give more rapid selection in this scenario (Figure 1K), the increase in resistance does not outweigh the increased control offered by higher doses (Figure 1L).

**Figure 1.**
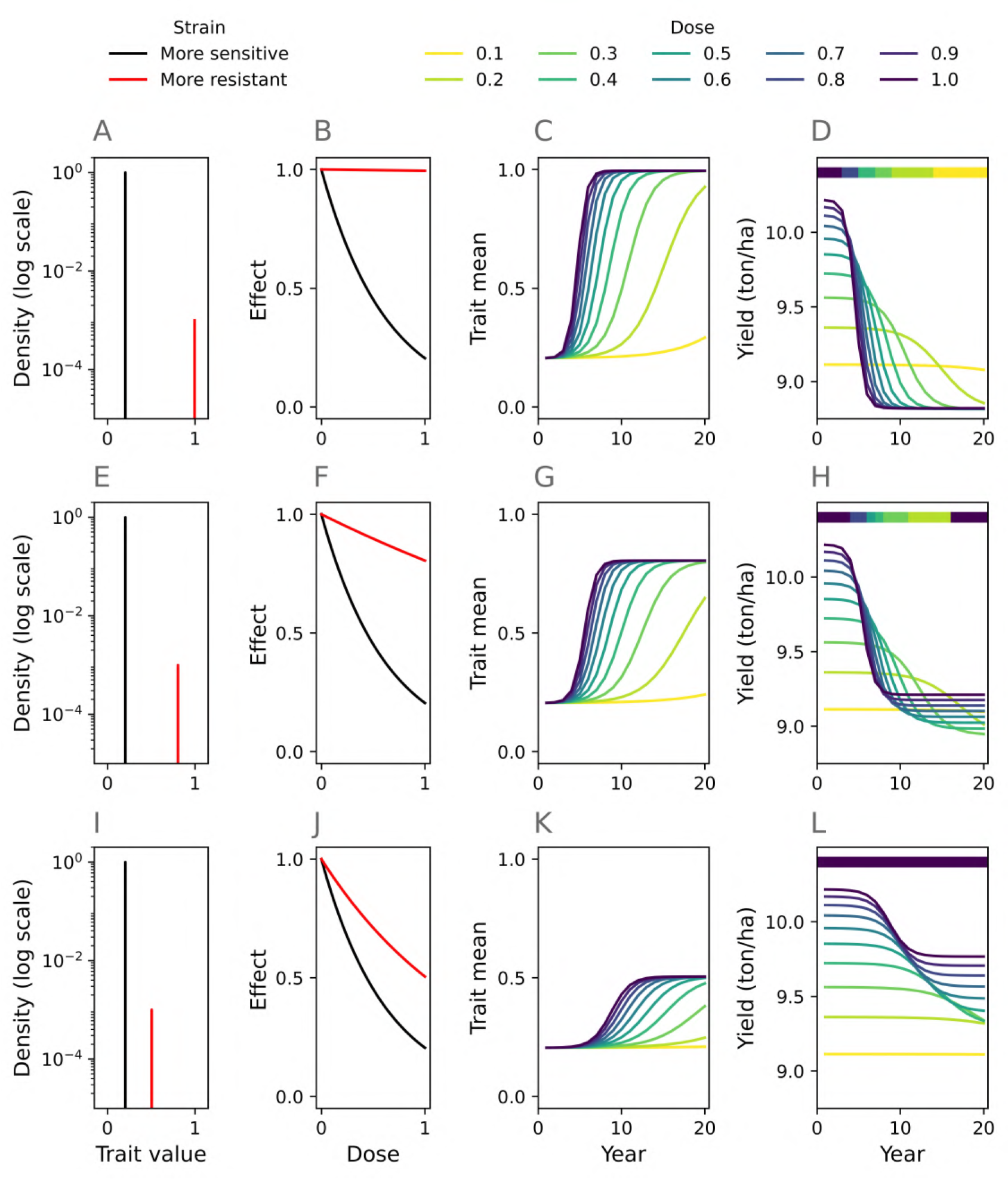
High doses can outperform low doses even in a qualitative resistance model. Each row corresponds to a single monogenic pathogen population evolving over 20 years. The left column shows the initial pathogen populations, each containing a sensitive and a resistant strain (**A**,**E**,**I**). Recall that *k* = 0 corresponds to a fully sensitive strain and *k* = 1 corresponds to a fully resistant strain. The middle-left column shows their corresponding dose response curves (**B**,**E**,**J**). The middle-right column shows how the mean trait value (Equation 9) in the population changes as resistance develops for various different fungicide doses applied each year (**C**,**G**,**K**). The right column shows how the yield varies as resistance develops for the same dose choices (**D**,**H**,**L**). The colourbars at the top of panels **D**,**H**,**L** show which dose is optimal in each year. The top row (**A**-**D**) shows an example where lower doses are best for control/yield after the first few years. In the middle row (**E**-**H**) the optimal dose in any given year depends on how long it has been since the strategy began – in the very short and very long term high doses are best but low doses are best in the intermediate period. In the bottom row (**I**-**L**) higher doses are best in all years. **Parameter values**. Resistant strain *k* value: **A**: 0.995; **E**: 0.805; **I**: 0.505. Sensitive stain *k* value: **A**,**E**,**I**: 0.205. Resistant strain initial density: 10^−3^. Mutation proportion: 0. Fungicide asymptote: 1.

To systematically explore when partial resistance affects optimality of high doses, we run a scan over the possible *k* values that the sensitive and resistant strains could take (Figure 2). For each year *Y* we plot the best dose, i.e. the one which gives the highest yield in year *Y* given the use of that dose in years 1, 2, … , *Y*. The doses considered are 0.1, 0.2, … 1. In Figure 2 we consider *Y* = 1, 5, 10, 15, 20.

**Figure 2.**
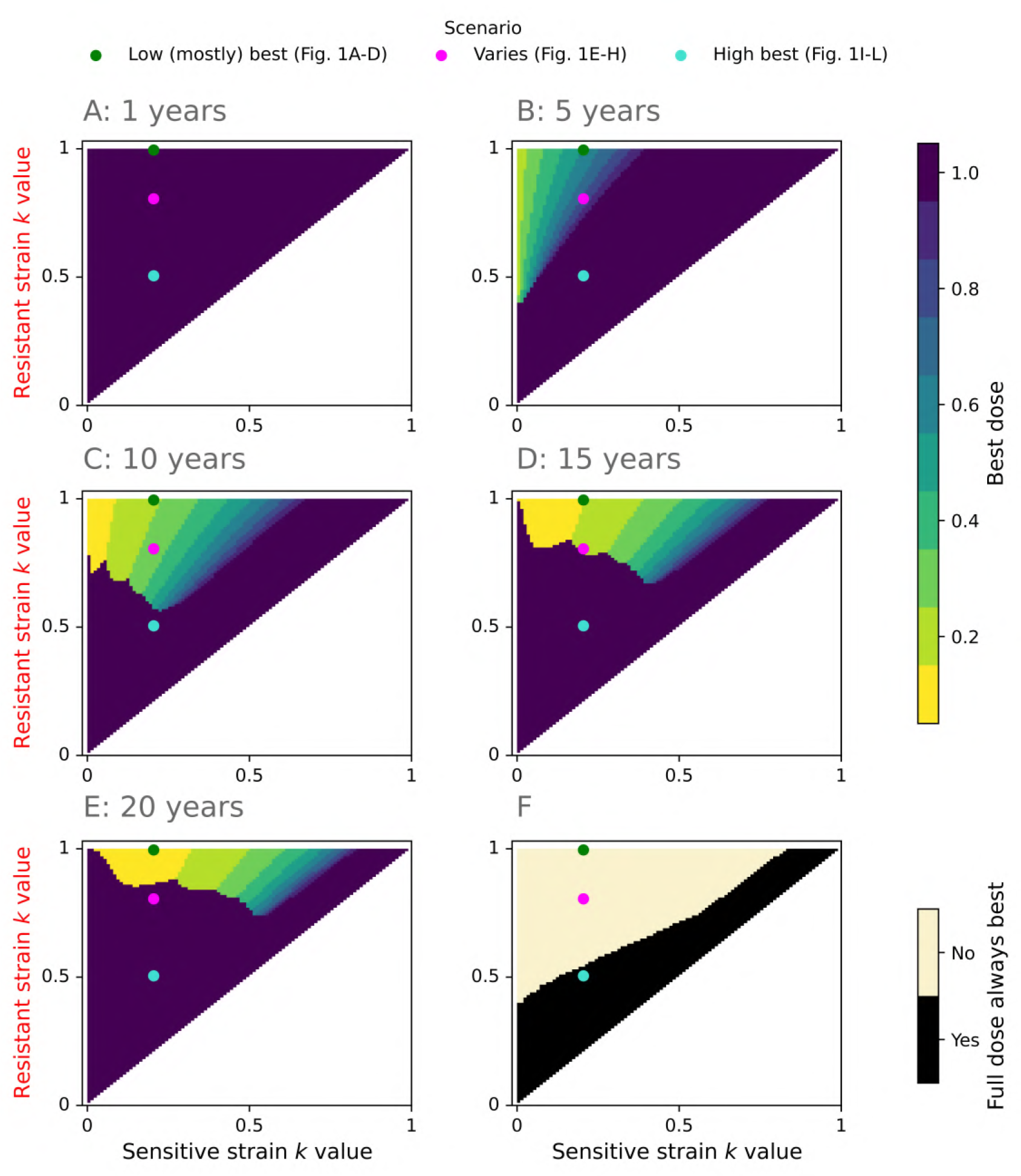
When do high doses outperform low doses in the qualitative resistance model? The resistant strain was constrained to take a higher *k* value than the sensitive strain, leading to the upper left triangle in each heatmap (recall that *k* = 0 is fully sensitive and *k* = 1 is fully resistant). We denote the three scenarios from Figure 1 by the three dots: green relates to Figure 1A-D; pink relates to Figure 1E-H; turquoise relates to Figure 1I-L. Yellow/green regions denote where using the low dose every year leads to a better yield in that year than using full dose every year, and (dark) purple where full dose is better. For some pairs of strains, high doses always give the best yield (**F**). **Parameter values**. Resistant strain initial density: 10^−3^. Mutation proportion: 0. Fungicide asymptote: 1.

In year 1, high doses are best for all strain pairings since they offer better control and the resistant frequency is low (Figure 2A). As time progresses, the best dose depends on the *k* values of the resistant and sensitive strains (Figure 2B-E).

The three scenarios from Figure 1 are shown in Figure 2 by the green, pink and turquoise dots. The green dot shows when the resistant strain is fully resistant (i.e. Figure 1A-D), but the pink (Figure 1E-H) and turquoise (Figure 1I-L) show when the resistance is partial (weaker in the turquoise dot example). In the green dot example (*k*_*s*_ = 0.205, *k*_*r*_ = 0.995), low doses give higher yields for all years from year 5 onwards (Figure 2B-E), although note that full dose is better in year 1 (Figure 2A). In the pink dot example (*k*_*s*_ = 0.205, *k*_*r*_ = 0.805), full dose gives higher yields initially and after many years (Figure 2A,E), but the lower dose is better in the interim period (Figure 2B-D). This is because full dose is better when the population is (almost) completely sensitive or completely resistant, but in the interim period the resistant proportion increases more rapidly when full dose is used. If the resistant strain is only weakly resistant (turquoise dot; *k*_*s*_ = 0.205, *k*_*r*_ = 0.505), then full dose always performs better or equal to the lower dose (Figure 2A-F).

When the resistant strain is very close in *k* value to the sensitive strain, high doses are always better since in this case the increased selection for resistance has a smaller effect than the increased control offered by a high dose (Figure 2F; pairs of values of *k* which are nearly equal are just above the line *k*_*s*_ = *k*_*r*_, which is shaded in black). When resistance is complete (i.e. *k*_*r*_ = 1), and the sensitive strain is highly sensitive (*k*_*s*_ ≈ 0), the low dose is better by year 5 (Figure 2B). In later years (Figure 2C,D,E), the population is essentially completely resistant, so full dose and the low dose both perform badly, offering very little control.

When the resistant strain is partially resistant (*k*_*r*_ < 1), then once enough time has passed that (virtually) the entire population is resistant, the full dose strategy outperforms the low dose strategy (since higher doses can control the partially resistant strain better). This means that after sufficiently many years, high doses will be best if resistance is partial (*k*_*r*_ < 1). However, the time taken to reach this entirely resistant stage is shorter with high doses due to increased selection from the full dose strategy, and the yield obtained when the population reaches this point may be unacceptably low. Depending on the two *k* values, there may be an interim period where low doses are better than high (yellow/green in Figure 2).

### When are high doses best in the quantitative resistance model?

We now explore which doses give the best yield over time in the quantitative resistance model. Understanding the behaviour of the qualitative resistance model is useful since it explains the behaviour of pairs of strains taking different *k* values (Figures 1, 2). The quantitative resistance model is more complex, with many different strains all taking different *k* values and all competing with each other.

We present the results of the parameter scan, which involved an ensemble of 10,000 model runs. For any given model run (e.g. Figure 3A), we can find the best dose in each year by comparing yield in that year for each of the 10 different doses (0.1, 0.2, … 1) tested. In the example shown, initially high doses are best but in later years lower doses are favoured (Figure 3A). This behaviour occurred in some model runs, but for many full dose was always favoured (Figure 3B). After 10 years, lower doses tend to be best for lower decay rates and higher fungicide asymptote parameters, corresponding to higher efficacy fungicides which remain at high concentrations for longer (Figure 3C). For a given mean curvature value, a higher ‘rate’ parameter (*b*) means decreased variance in the distribution of curvature values. Note that the rate parameter *b* influences the shape of the initial pathogen trait distribution and despite its name it is not an infection rate or a parameter with any time-dependence. Higher mean curvature values and lower rate parameters tend to incentivise lower doses (Figure 3D). The effect of the gamma distribution parameters on the resulting trait distribution is shown in Appendix 3 Figures 1, 2. The lower the mean effect at full dose, corresponding to higher efficacy fungicides which suppress the disease more strongly, the more likely that lower doses will be best (Figure 3E). Higher mutation scales tend to correlate with lower doses being preferable, but the mutation proportion is less important (Figure 3F).

**Figure 3.**
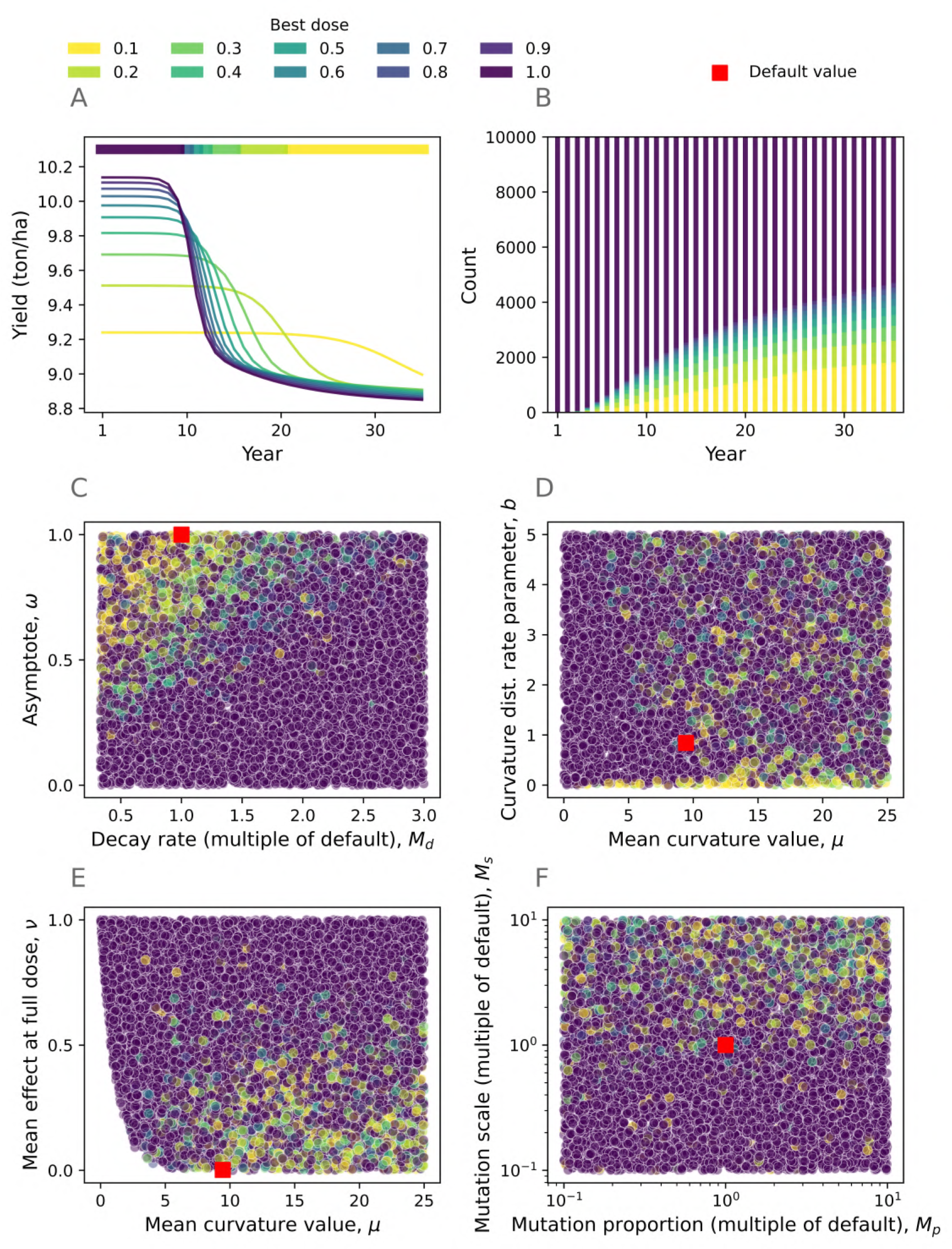
High doses can outperform low doses in the quantitative resistance model. Panel **A** shows one example model run from the ensemble. The ‘best dose’ in any given year is the dose which gives the highest yield in that year, as shown by the colourbar at the top of panel **A**. Panel **B** shows how frequently higher doses outperformed low doses depending on the timescale of interest. Initially, high dose is invariably best since the population is still highly sensitive, but after some time low doses are better for some of the parameter combinations generated in the scan. Panels **C**-**F** explore which parameters led to low doses outperforming high doses, focusing on year 10 only (arbitrarily chosen). This helps us understand which parameters drive the effect. Higher efficacy fungicides, i.e. with a high dose-response asymptote (**C**), a low decay rate (**C**), a high mean curvature value (**E**) or a low average maximum effect (**E**) require lower doses. Note that the mean effect at full dose *ν* is a function of *μ*, *b* and *ω* as described in Equations 10 and 11, which results in the shape of the space in panel **E**. Populations with greater mutation scales require lower doses **F**, but the mutation proportion is less important. There is less of a clear pattern in **D**, although low doses seem better for high values of mean curvature *μ* and for low values of the rate parameter *b*. Default values as in ***Taylor and Cunniffe*** (***2022a***) are denoted by red squares. Parameter values for panel **A** are in Appendix 1 Table 2, and a plot of how the pathogen distributions from panel **A** change with different doses is in Appendix 3 Figure 3.

Shapley values sum to the difference between the baseline/expected model output and the current model output for a particular prediction being explained. Each variable in each year of each model run has a single Shapley value. For another single model run, we can see how the yield varies for different doses (Figure 4A). The waterfall plots in Figure 4B,D each show a single prediction from the model, corresponding to the first year of this model run and the 20th year respectively. The gradient boosted model uses the explanatory features (Table 3) to predict which dose gives the optimal yield in each year. The Shapley values explain how the prediction differs from the expected best dose of 0.811 (averaged across the entire dataset). Features are added in one at a time until we reach the model output of 1.003 in the first year (i.e. best dose is full dose, plus a small model error). Note that the model output is a continuous variable, so in general it won’t equal a multiple of 0.1. The main contribution in year 1 is from the year variable *Y* - the best dose to use in the first year is full dose since it provides improved control in that year and resistance is minimal. For this model run, we plot the impact of the year variable Figure 4C. Here we plot the feature value (year, *x* axis) against the corresponding SHAP value (*y* axis). As time progresses the SHAP value in each year of this model run decreases, meaning that higher doses are best in early years but lower doses are best if longer timescales are considered. In the 20th year, mutation scale (*M*_*s*_) and mean effect at full dose (*ν*) are the variables with the largest contributions (Figure 4D). The best dose predicted by the gradient-boosted trees model for the 20th year is 0.231. In the actual model run (**A**), 0.2 was the best dose in year 20 and 0.3 was the best dose in year 19.

**Figure 4.**
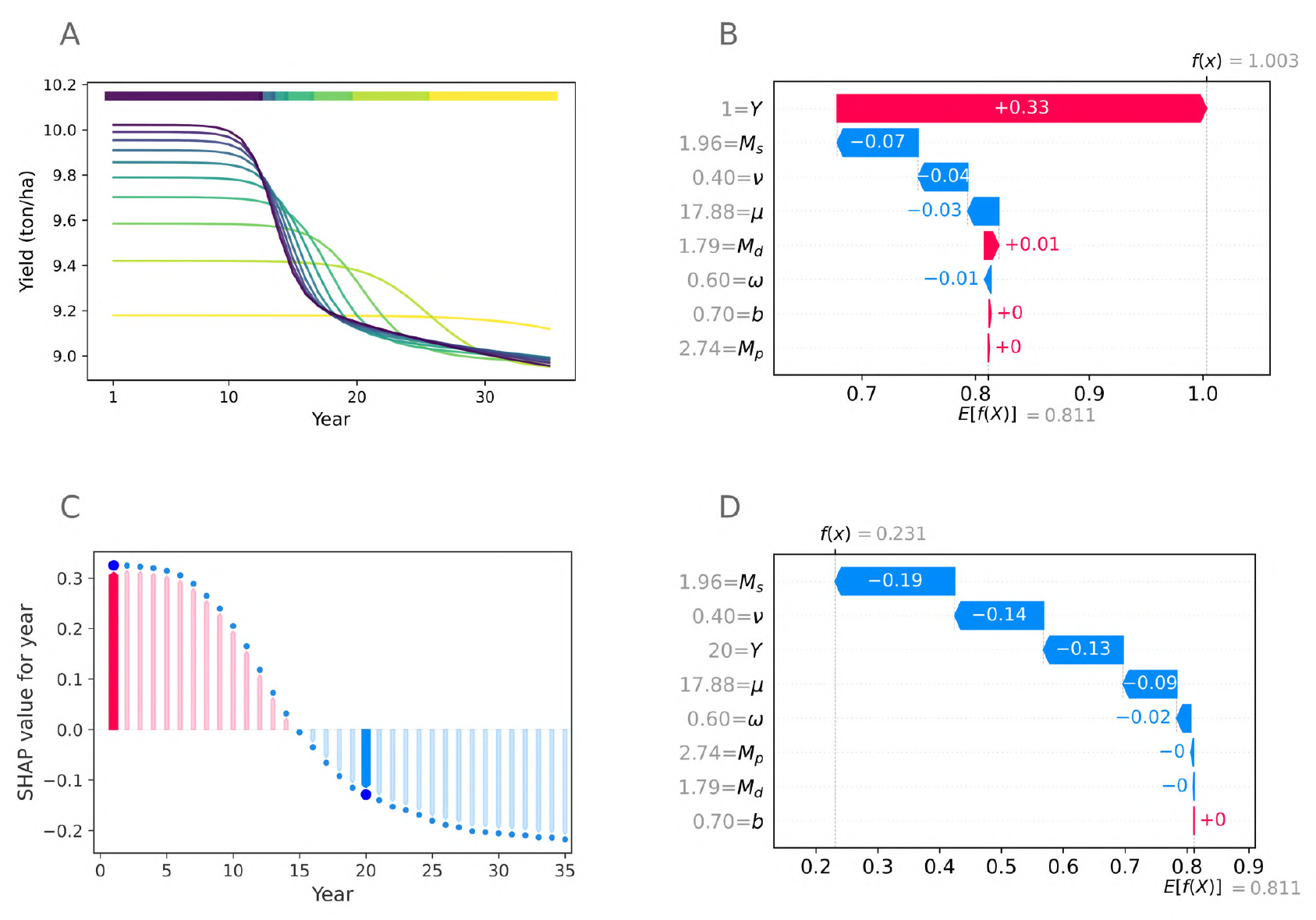
Introducing SHAP values. The best dose in any given year varies in this single model run from the ensemble of 10,000 (**A**). Initially full dose is best, but in later years lower doses are optimal. Panel **B** shows a ‘waterfall plot’ explaining a single model prediction corresponding to the first year from this model run. In this example the year, *Y*, is the most important feature and the predicted best dose is 1.003 (i.e. full dose with a small model error). See Table 3 for an explanation of the features and their ranges. Panel **C** shows, for this model run, how the year features influences the model output. Higher doses are better in early years, which is reflected in positive contributions when the year is small and negative contributions when the year is larger than 15. The first year and 20th years are highlighted since these are the data points shown in more detail in **B** and **D** respectively. Panel **D** shows a waterfall plot for the 20th year from this model run. The predicted best dose is 0.231 and the most important feature is the mutation scale *M*_*s*_. Parameter values for panel **A** are in Appendix 1 Table 2.

Feature importance is shown in Figure 5A, and summarises the influence of each parameter on the optimal dose. This is measured in terms of the mean absolute SHAP value over the entire dataset. Figure 5B shows the relationship between the features and the SHAP value in more detail. Each year of every model run is represented by a single point for each feature. Higher values for each feature are shown in blue, and lower values in pink. The larger the absolute SHAP value, the bigger the impact of that point on the model output. The most important variables are the mutation scale (*M*_*s*_), the year (*Y*) and the mean effect of fungicide at full dose (*ν*) (Table 3).

**Figure 5.**
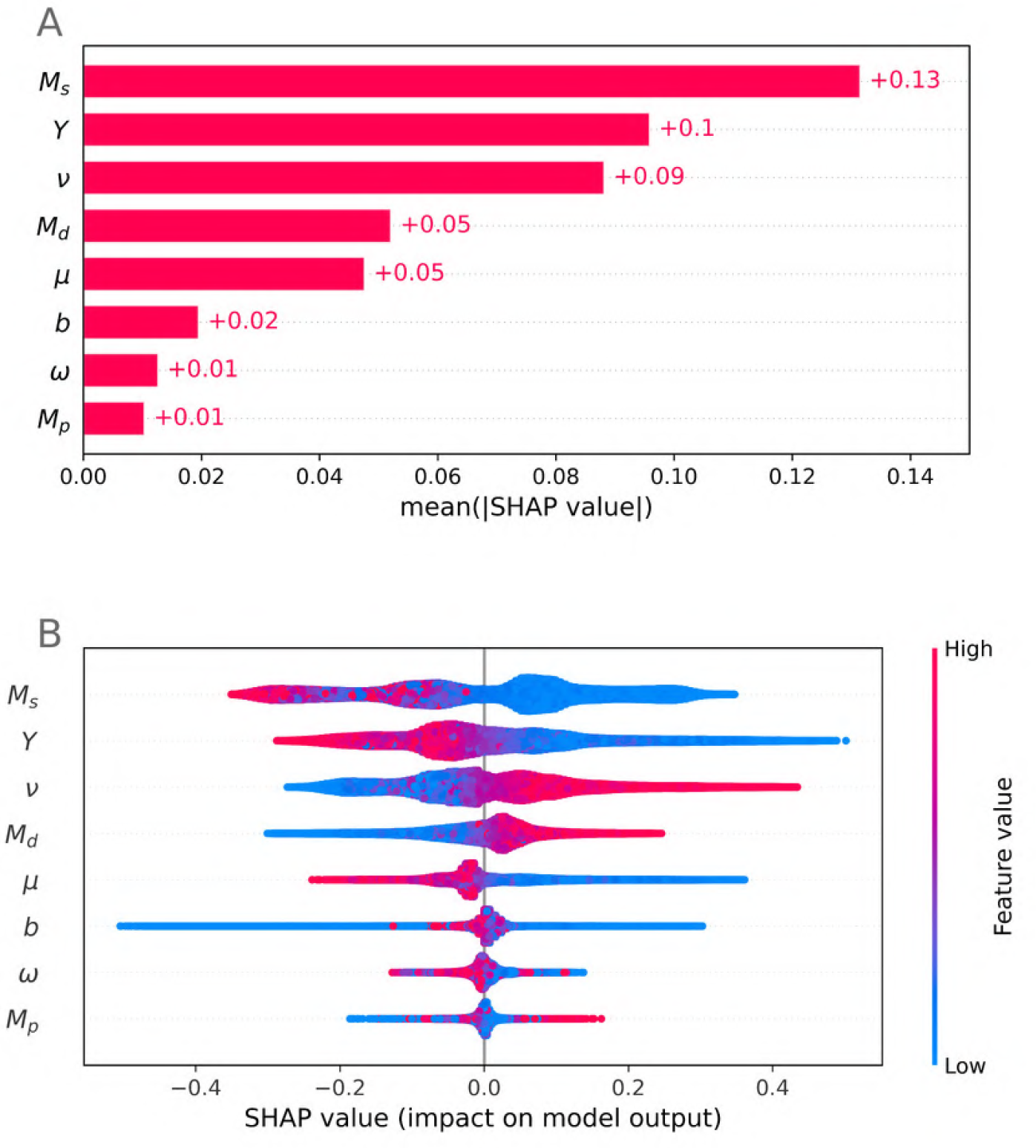
Which variables have the greatest impact on determining the optimal dose? Panel **A** shows feature importance, measured in terms of the mean absolute SHAP value over all given samples. Mutation scale *M*_*s*_, year *Y* and fungicide efficacy (mean maximum effect *ν*) are the most important features. Panel **B** shows the effect of the most important variables on the model output across the scan. Each point corresponds to a single year from a single model run. Blue points for any given variable correspond to low values for that variable, and similarly pink points represent high values for that variable. Their impact on the model output is shown on the *x* axis. The density of points is represented by their height (comparable to a violin plot). Panel **B** a more detailed explanation of the impact of each variable on the model output than panel **A**.

To further understand the effect of each variable, we plot the SHAP value against each parameter value showing the results for all members of our ensemble (Figure 6). This is the same style of plot as Figure 4C, but we show all data from all model runs rather than only those from a single model run. In all panels we use colour to show the interaction with the year (NB the colours in Figure 6A do not relate to an interaction since the variable plotted in Figure 6A is year). To further understand the effect of the model parameters without the year interaction, we fit a model to the data from year 10 only and performed similar analysis in Appendix 4, and found that the order of feature importance remains the same, apart from the mean effect at full dose and decay rate features swapping importance order from 2nd and 3rd most important to 3rd and 2nd most important.

**Figure 6.**
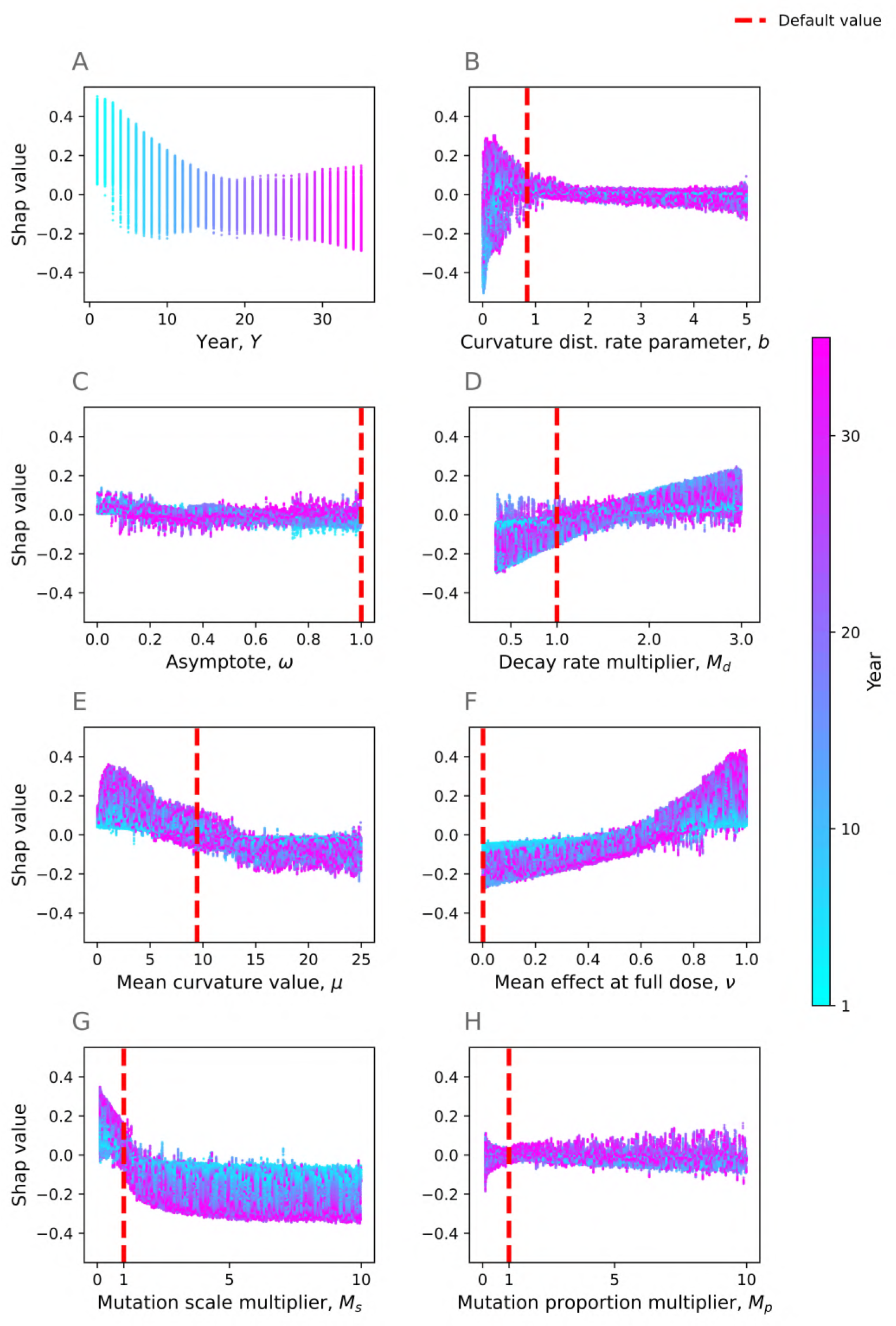
Which variables correlate with optimality of high doses? Here we show the effect of many of the model variables on the output. Each point corresponds to a single year from a single model run in the ensemble. Each *x*-axis value corresponds to a variable value, and each *y* axis is the SHAP value for that feature, corresponding to how much that feature contributes to the model output for that specific prediction. A high SHAP value corresponds to that variable changing the output to a the optimal dose being higher. Default model values as fitted in ***Taylor and Cunniffe*** (***2022a***) are shown by the red dotted lines. The colours help indicate interactions between each feature and the feature ‘year’ (*Y*). The model predicts higher doses for decreased year (**A**) and decreased fungicide efficacies, i.e. higher decay rates (**D**), lower mean curvatures (**E**), and lower mean effect at full dose for the initial pathogen distribution (**F**). However, the model predicts higher doses for increased decay rates (**D**), mutation scales (**G**) and proportions (**H**). The curvature rate parameter *b* (**B**) and mutation proportion multiplier (**H**) features have a smaller effect, although very small values of *b* have a more significant effect. This is because they relate to higher pathogen density at very low and very high *k* values (see Appendix 3). The asymptote variable *ω* also has a relatively small effect (**C**), most likely because its impact on the model is via its impact on the mean effect at full dose *ν*.

The predicted best dose decreases strongly with the year and mutation scale (Figure 6A,G) but increases strongly with the decay rate and the mean effect at full dose (Figure 6D,F). This matches the result shown in Figure 5A showing the most important 4 features as summarised by average absolute Shapley value to be mutation scale, year, mean effect at full dose, and decay rate. Higher mean curvature values *μ*, (corresponding to higher efficacy fungicides) mostly correspond to lower doses being optimal (Figure 6E). The asymptote feature, *ω*, has quite a small impact on the model (Figure 6C), most likely since its impact is essentially summarised by the mean effect at full dose feature *ν*. The curvature rate parameter *b* has a small effect when it is greater than 1 but a large, though unpredictable effect when it is smaller than 1 (Figure 6B). This is because the effect of increased variance in the pathogen distribution interacts strongly with other variables, and leads to large changes in the initial pathogen distribution with high density near *k* = 0 and *k* = 1 (Appendix 3 Figures 1). The mutation proportion multiplier has a smaller effect than the mutation scale (Figure 6H).

Many of the variables interact strongly with year. Notably the mean curvature value (*μ*, Figure 6E), the mean effect at full dose (*ν*, Figure 6F) and the mutation scale (*M*_*s*_, Figure 6G) have an exaggerated effect on the model in later years but a much smaller effect in early years. This is because high doses are favoured in early years but there is bigger variation in possible outcomes in later years depending on whether low doses begin to outperform high doses.

#### Biological interpretation

The parameter scan suggests that higher efficacy fungicides tend to need lower doses for optimality, after the first few years in which high doses are always best (Figures 3,5,6). This is because the selection for resistant strains is stronger – if the initial mean effect at full dose is very low, corresponding to a higher efficacy fungicide, then the difference in growth rates between sensitive and resistant strains is larger so resistance develops more quickly (according to the so-called governing principles (***van den Bosch et al., 2014***)) and low doses become preferable to higher doses after a small number of years (Figure 7A,B). Making a comparison with the monogenic case (Figure 2), lower efficacy fungicides have the sensitive strain(s) at a relatively high *k* value(s), away from *k* = 0, which tends to incentivise higher doses unless the resistant strain is close to *k* = 1. This means that in most cases lower efficacy fungicides correlate with higher doses being optimal (Figure 7C,D). However, if the mutation scale is very high then the pathogen population reaches higher density near *k* = 1 quicker (Figure 7E,F), unless doses are lowered greatly which reduces selection. Note that the increasing the mutation proportion has a smaller effect than increasing the mutation scale (Figure 7G,H).

**Figure 7.**
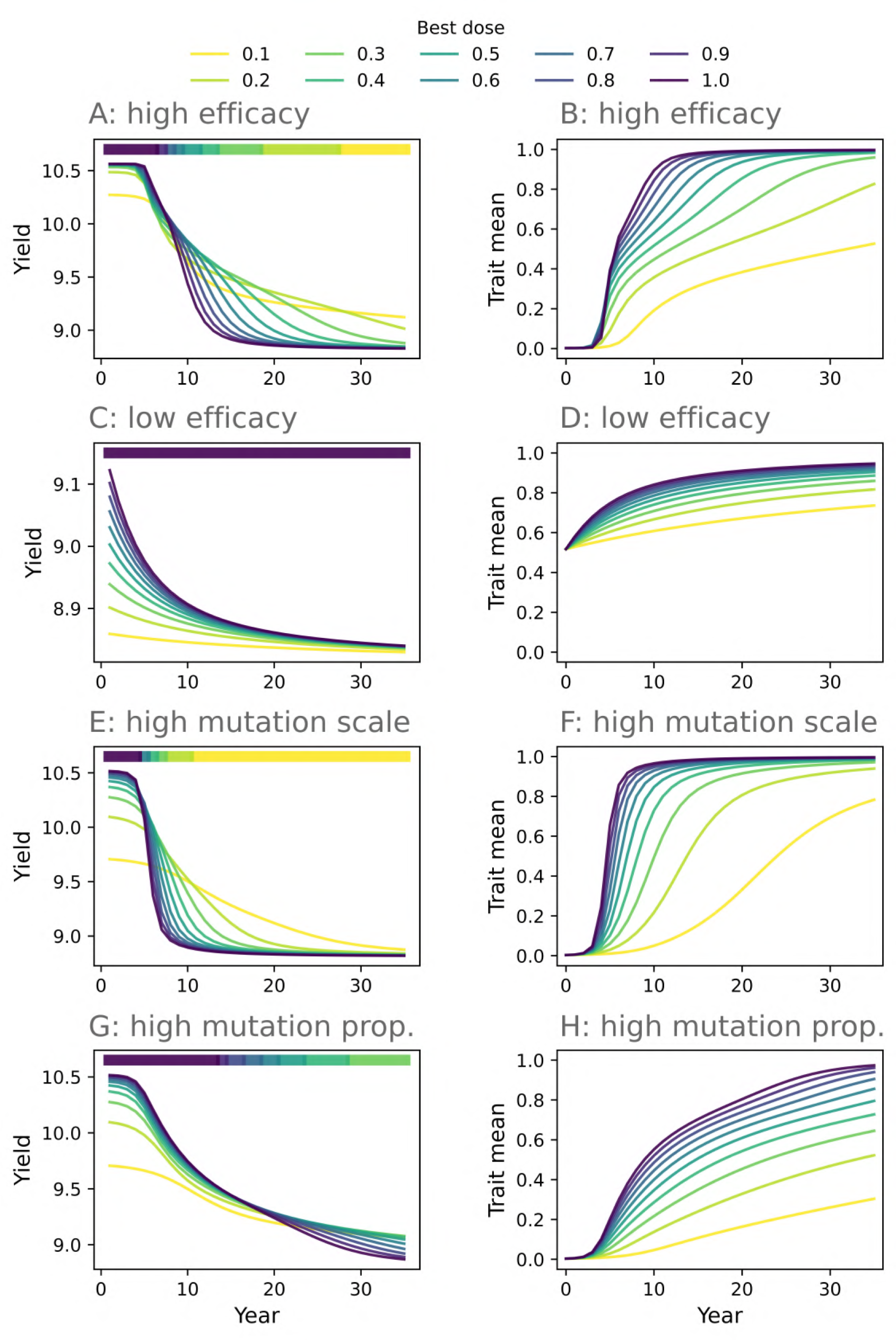
Why does fungicide efficacy and pathogen mutation affect the dose recommendation? High doses of high efficacy fungicides select strongly for resistance, leading to lower doses being optimal after only a few years (**A**, **B**). However, higher doses of lower efficacy fungicides can provide more effective control without selecting too rapidly for resistance (**C**, **D**). Large mutation scales lead to rapid resistance development when high doses are used, incentivising lower doses (**E**, **F**). Large mutation proportions do not lead to as rapid resistance development if the mutation scale is not particularly large (**G**, **H**). In this scenario it is possible for high doses to be best for many years. **Parameter values**: see Appendix 1 Table 3.

In all examples shown, higher doses give increased selection for resistance (Figure 7B,D,F,H). However, this does not guarantee reduced control and correspondingly reduced yield if the difference in selection is small relative to the increase in control offered by a higher dose (compare Figure 7A,E,G with Figure 7C).

## Discussion

Fungicide dose choice can be important in determining how quickly resistance develops (***van den Bosch et al., 2011***). Although the existing literature suggests that higher doses contribute to increased selection (***van den Bosch et al., 2011***, ***2014***; ***Hobbelen et al., 2013***; ***Elderfield et al., 2018***; ***Taylor and Cunniffe, 2022b***), many studies on optimal dosage neglect partial resistance, and to the best of our knowledge no modelling study considers the case of quantitative resistance. We show that although higher doses can often lead to increased selection for resistance in these scenarios, the control offered by higher doses may still outperform that offered by lower doses, both in the case of partial qualitative resistance (Figures 1, 2) and in quantitative resistance (Figure 3, 4).

In the early years of a fungicide programme, high doses always give the best control. However, in later years the optimal dose choice depends on model parameters (Figures 4, 5, 6). High fungicide efficacies often incentivise lower doses. This can be in terms of the decay rate of the fungicide itself, or in its interaction with the pathogen via the dose response curve: the asymptote, mean curvature or (initial) maximum effect at full dose can be implicated - see Figures 6C,D,E,F. This is because the difference in infection rates between resistant and sensitive strains is greater with high efficacy fungicides, leading to greater selection (Figure 7A-D) according to the governing principles as originally developed for monogenic resistance (***van den Bosch et al., 2014***). In Figure 7A,B the rapid selection for resistant strains outweighs the benefit in control offered by increased dose, whereas in Figure 7C,D the difference between the resistant and sensitive strains is smaller, so selection is less exaggerated and the resistance does not outweigh the benefit to control offered by increasing fungicide dose.

Increased pathogen mutation scales also incentivise lower doses (Figure 6G). This is because increased mutation scales can rapidly lead to higher densities of highly resistant strains. These strains are selected for more strongly if doses are higher, so low doses are best in this case (Figure 7E,F). However, increased mutation proportions may not have the same effect unless mutation scales are also increased (Figure 6H, Figure 7G,H).

The results characterised the optimal dose solely in terms of instantaneous yield, i.e. the yield in any given year, rather than the total yield up until that point. Using instantaneous yield ignores the magnitude of the difference between the optimal yield and sub-optimal yields. This difference may be much larger in some years than others. To explore the effect of optimising for the total yield rather than instantaneous yield, we ran a similar analysis combining a gradient-boosted trees model with Shapley values (Appendix 5). The results were broadly similar, although the initial (sometimes large) benefit to using high doses meant that there were more parameter combinations which favoured high doses (Appendix 5 Figures 1, 2). The most important explanatory variables were similar, although year became less important than the effect of the decay rate multiplier *M*_*d*_ (Appendix 5 Figures 3, 4).

In ***Mikaberidze et al***. (***2017***) dose-convergence (***van den Bosch et al., 2011***) is used to explain the potential for higher doses to delay emergence of resistance when resistance is monogenic and partial (and affects the ‘slope’ of the dose response rather than the maximum effect, i.e. ‘Type 2’ partial resistance). Dose-convergence means the difference in growth rates between sensitive and resistant strains is smaller at high doses than at lower doses. For some models this can theoretically lead to higher doses exerting lower selection pressure on resistant strains. In this work, resistance is characterised in terms of the curvature parameter, making resistance in the model analogous to Type 2 partial resistance. However, we use a different dose-response curve parameterisation and assume that the fungicide concentration decays exponentially (following ***Hobbelen et al***. (***2013***); ***Elderfield et al***. (***2018***); ***Taylor and Cunniffe*** (***2022b***); ***Caffi and Rossi*** (***2018***)) rather than remaining constant. The decay in fungicide concentration largely negates the dose-convergence argument (ignoring density dependent effects). This is because even if selection is lower when a high dose is applied, there will be stronger selection as the chemical decays because the concentration passes through all lower doses. This means that the selection for resistant strains is greater for higher doses, even in the quantitative resistance case. However, dose-convergence may limit the extent to which selection increases as dose increases. This is illustrated by the model in Appendix 6, which shows that higher doses dominate less frequently when resistance is characterised by the fungicide asymptote parameter (analogous to partial resistance Type 1). Unlike ***Mikaberidze et al***. (***2017***), our focus was on the yield obtained from using different doses, rather than the effect of dose choice on emergence of resistance. This means we take into account the benefit in control from using a higher dose. In the quantitative resistance case the optimal yield was from full dose for over 5,000 model runs (from the ensemble of 10,000 in the randomisation scan) in every single model year (Figure 3B).

***Hobbelen et al***. (***2013***) show that, for mixtures of fungicides to which resistance is qualitative and partial, substantially higher than minimal doses maximised lifetime yield (time for which acceptably high yields can be maintained), particularly if the partial resistance was Type 2 partial resistance. Conversely, ***van den Bosch et al***. (***2011***) state that dose-convergence is unlikely to be relevant to single-site fungicides (i.e. the monogenic/qualitative resistance case), since dose-convergence is unlikely to occur within a legal range of doses when there is a single genetic change which confers a high level of resistance, However, they suggest it may be relevant in for azole fungicides (i.e. quantitative resistance). Our results suggest that high doses still tend to lead to greater levels of resistance in the quantitative resistance case, again due to exponential fungicide decay negating the dose-convergence argument. Nevertheless, for both quantitative and partial qualitative resistance, this increased selection may not be sufficient to make low doses optimal in terms of yield, since the control offered by increasing dose may outweigh the increased selection if resistance is not complete.

Various modelling assumptions were made when deriving the model. Fitness costs are neglected, which may have a significant impact (***Hawkins and Fraaije, 2018***). However, ***Mikaberidze et al.*** (***2017***) did not find fitness cost to have any major effect on their results on emergence time for fungicide resistance. Although the current form of our model does not account for Type 1 partial resistance (i.e. where resistant strains in the population differ according to the maximum effect of the fungicide), we note that the primary difference between partial resistance Type 1 and Type 2 in behaviour stemmed from dose-convergence which is less significant in our model due to assuming fungicide concentration decays rather than remains constant. For simplicity, spatial effects are ignored, although future work may consider them given the potential for fungicide strategies based on spatial risk (***Liu et al., 2017***). We also neglect to model Septoria’s latent period (***Fones and Gurr, 2015***) for simplicity. Although we consider emergence of new strains in the population via mutation, there is no demographic stochasticity, meaning that strains at low densities have zero probability of dying out. Demographic stochasticity could be important to include since one of the existing arguments for higher doses is delaying emergence of resistance via reducing the population size and therefore reducing the absolute number of mutant offspring (***van den Bosch et al., 2011***).

Although the model in ***Taylor and Cunniffe*** (***2022a***) is capable of addressing disease resistant cultivar control, for simplicity we omitted it here. However, it would be interesting to explore whether introducing cultivar control had any effect on the optimal dose. In early years lower doses could be used whilst retaining acceptable control if disease-resistant cultivars were used, but equally higher doses could offer increased protection to the host, delaying the breakdown of cultivar control (***Carolan et al., 2017***). Much of the fungicide resistance modelling literature is concerned with mixtures containing fungicides challenged by qualitative resistance (***Hobbelen et al., 2013***; ***Elderfield et al., 2018***; ***Taylor and Cunniffe, 2022b***). The model could be extended to help finding optimal doses depending on initial conditions and the dose-responses of the fungicides in the mixture in the quantitative resistance case, or the case where one fungicide faced qualitative resistance and the mixing partner faced quantitative resistance. Finally, it would be interesting to explore time-variable tactics. For example, in earlier years acceptable control could be obtained using low doses, and dose only increased once necessary to achieved desired levels of control. An economic analysis accounting for fungicide costs would also be interesting (***Alves et al., 2021***; ***van den Bosch et al., 2020***). This approach would help us understand how to delay resistance development whilst maintaining good crop and economic yields for as long as possible. Nevertheless, by showing how both high and low doses can lead to optimal yields over different timescales, as well as understanding the drivers of this behaviour, we have shown how optimal dose choices for quantitative and/or partial qualitative resistance may frequently differ from optimal choices when resistance is qualitative and complete.

## Availability of code online

An implementation of the model in the freely-available programming language Python (Python Software Foundation, available at python.org) is online at github.com/nt409/polygenic2.

## Acknowledgements

NPT acknowledges the Biotechnology and Biological Sciences Research Council of the United Kingdom (BBSRC) for support via a University of Cambridge DTP PhD studentship (Project Reference 2119688).

## Appendix 1

### Parameter values

#### Default parameter values

The default parameter values, as fitted in ***Taylor and Cunniffe*** (***2022a***).

**Appendix 1 Table 1.** Default parameter values used in the model (to 3 significant figures). Only the top 4 are actually used in each model run in the parameter scan, since we randomly sample values for the others.

| Parameter | Symbol | Value |
| --- | --- | --- |
| Initial inoculum | $I_0$ | 0.00986 |
| Infection rate (no control) | $\beta_0$ | 0.00787 |
| Host growth rate | $r$ | 0.0126 |
| Host initial area | $H_0$ | 0.0122 |
| Fungicide decay rate | $\Lambda$ | 0.00901 |
| Fungicide asymptote | $\omega$ | 1 |
| Pathogen gamma distribution mean | $\mu$ | 9.44 |
| Pathogen gamma distribution rate parameter | $b$ | 0.843 |
| Mutation proportion | $p_M$ | $1.23 \times 10^{-5}$ |
| Mutation scale | $\sigma^2$ | 0.00192 |

#### Parameter values, Figures 3A, 4A and Appendix 5 Figure 2A

The parameter values for the model runs in Figures 3A, 4A and Appendix 5 Figure 2A.

**Appendix 1 Table 2.** Parameters values used in Figures 3A, 4A and Appendix 5 Figure 2A (to 3 significant figures).

| Parameter | F3A | F4A | A5 F2A |
| --- | --- | --- | --- |
| $\mu$ | 21.8 | 17.9 | 19.4 |
| $b$ | 3.89 | 2.74 | 1.32 |
| $\omega$ | 0.979 | 0.603 | 0.456 |
| $M_d$ | 2.46 | 1.79 | 1.85 |
| $M_s$ | 3.64 | 1.96 | 1.72 |
| $M_p$ | 0.837 | 0.704 | 0.109 |

#### Parameter values, Figure 7

The parameter values for the model runs in Figure 7.

**Appendix 1 Table 3.** Parameters values used in Figure 7 (to 3 significant figures). All other parameters not quoted here were at default values. The bold values are the non-default values. Scenarios: **A/B**: high efficacy fungicide; **C/D**: low efficacy fungicide; **E/F**: high mutation scale (lower mutation proportion); **G/H**: high mutation proportion (lower mutation scale).

| Parameter | Panels A/B | Panels C/D | Panels E/F | Panels G/H |
| --- | --- | --- | --- | --- |
| $\mu$ | **12** | **1** | 9.44 | 9.44 |
| $\omega$ | 1 | **0.5** | 1 | 1 |
| $M_d$ | **0.35** | **2** | 1 | 1 |
| $M_s$ | 1 | 1 | **7.5** | **0.5** |
| $M_p$ | 1 | 1 | **0.5** | **7.5** |

## Appendix 2

### Gradient-boosted trees model

#### Hyperparameter optimisation

The XGBoost hyperparameters optimised.

**Appendix 2 Table 1.** XGBoost (***Chen and Guestrin, 2016***) hyperparameters tuned during model fitting. We used the open-source python package ‘optuna’ for the optimisation procedure.

| Parameter | Explanation | Range | Distribution |
| --- | --- | --- | --- |
| Max depth | Maximum depth of each tree | [3, 20] | Uniform (integer) |
| Number of estimators | Number of trees in the model | [10, 2000] | Log-uniform (integer) |
| Learning rate | Learning rate / shrinkage | $[10^{-3}, 0.3]$ | Log-uniform |
| Subsample | The fraction of observations randomly sampled for each tree | [0.5, 1] | Uniform |
| Column sample by tree | Subsample ratio of columns used when constructing each tree | [0.4, 1] | Uniform |

**Appendix 2 Table 2.** Optimal hyperparameter values for each of the models, quoted to 3 significant figures. The ‘default’ model is the model used in the main text, corresponding to instantaneous yield and partial resistance Type 2. The ‘Year 10’ model is the model fitted to the data from the 10th year of each model run only (Appendix 4). The ‘cumulative yield’ model is the model fitted to the cumulative yield rather than instantaneous yield data (Appendix 5). The ‘partial resistance Type 1’ (PR Type 1) model is the model fitted to the results of the scan when resistance is characterised in terms of the fungicide asymptote rather than curvature (Appendix 6). Note that the optimal hyperparameter values for both the default and partial resistance Type 1 models are identical. In all cases, the hyperparameters were obtained through randomly sampling 10,000 hyperparameter combinations from the distributions in Table 1. The best parameters were selected by choosing the combination which minimised the cross-validation score on the training dataset (see Table 3).

| Parameter | Default | Year 10 | Cumulative yield | PR Type 1 |
| --- | --- | --- | --- | --- |
| Max depth | 10 | 8 | 8 | 10 |
| Number of estimators | 1740 | 1129 | 824 | 1740 |
| Learning rate | 0.0199 | 0.0226 | 0.0383 | 0.0199 |
| Subsample | 0.547 | 0.618 | 0.815 | 0.547 |
| Column sample by tree | 0.888 | 0.863 | 0.973 | 0.888 |

#### Model performance

The model performance in the training and testing stages, in terms of root mean squared error.

**Appendix 2 Table 3.** Gradient-boosted trees model performance. The loss function in model training is the squared error; here we quote the root mean squared error (RMSE). The cross validation process trains the model on splits of 6400 model runs (train) : 1600 model runs (validate) from the 8000 model runs used for model validation. Each model run contains 35 data points (one per model year), meaning that the cross validation training data contained 6400 × 35 = 224,000 rows of data. For the final model testing we used all 8000 training model runs to train the model before testing using the remaining 2000 previously unseen model runs. As is to be expected, the model performance is better on the training dataset. The model performance on the completely unseen data is comparable to the score obtained during cross-validation for the optimal hyperparameters for each of the models, meaning that the optimal model in each case generalises well to unseen data / does not degrade significantly. Values quoted to 3 significant figures.

| Dataset | Default | Year 10 | Cumulative<br>yield | PR Type 1 |
| --- | --- | --- | --- | --- |
| Training dataset, 5-fold cross validation (default hyperparameters) | 0.132 | 0.108 | 0.0617 | 0.150 |
| Training dataset, 5-fold cross validation (optimal hyperparameters) | 0.109 | 0.0947 | 0.0493 | 0.117 |
| Full training dataset (having trained on full training dataset) | 0.0373 | 0.00804 | 0.0164 | 0.0397 |
| Test dataset (having trained on full training dataset) | 0.106 | 0.0967 | 0.0437 | 0.116 |

Model performance on completely unseen data (the test set).

**Appendix 2 Figure 1.**
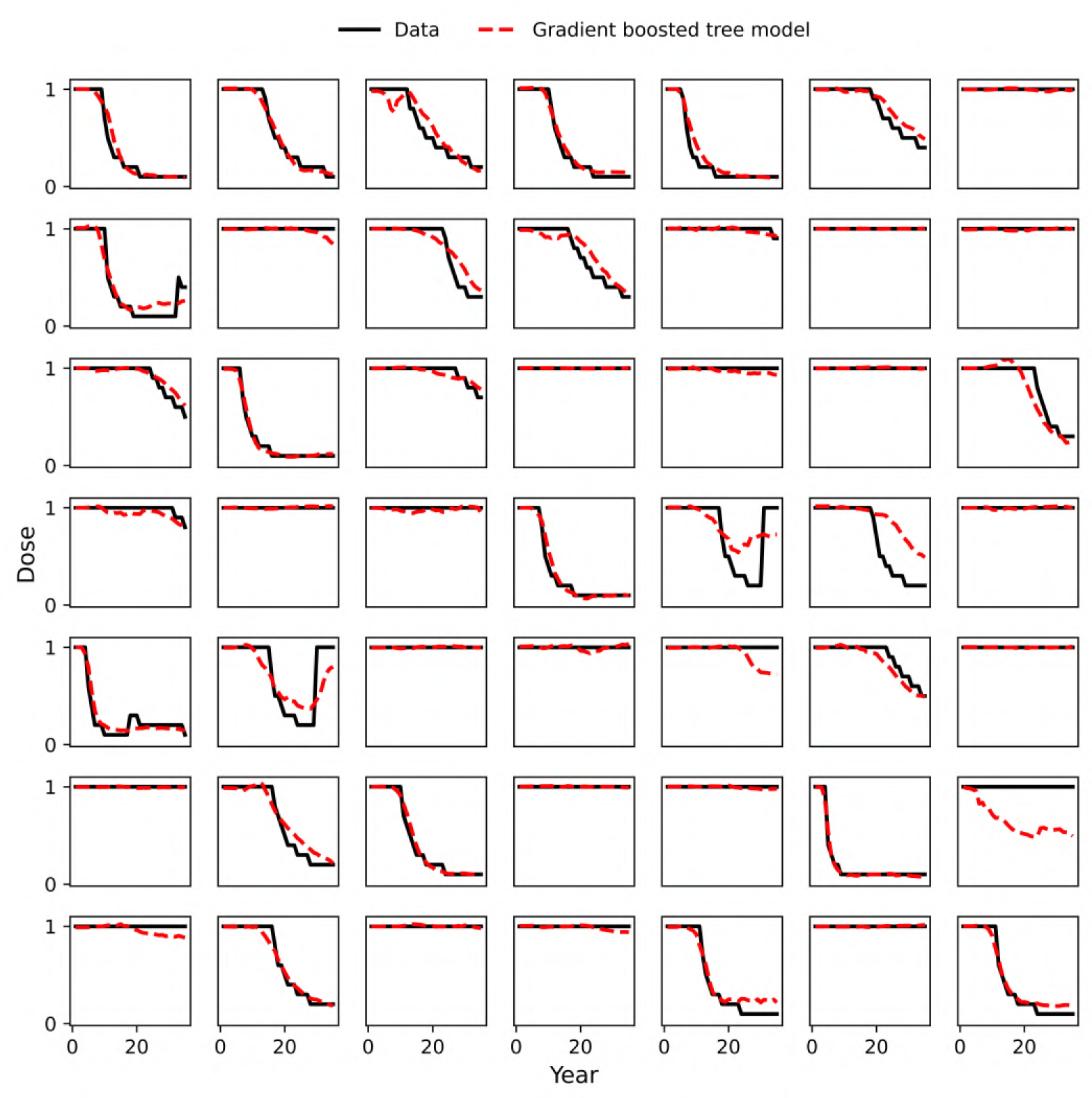
Gradient-boosted trees model performance on the test set – example model runs. Model performance on the first 49 of the 2000 model runs in the test set. The hyperparameters were optimised and model trained on the first 8000 runs in the ensemble of 10,000, meaning that the 2000 runs in the test set represented completely unseen data. For the vast majority of model runs the model performs well, accurately predicting when/if the optimal dose deviates from full dose. A larger training set might help to further reduce the small number of runs where the model performs poorly, which may occur if a run in the test set behaves differently (i.e. the optimal dose is very different) to a run in the training set with similar feature values.

**Appendix 2 Figure 2.**
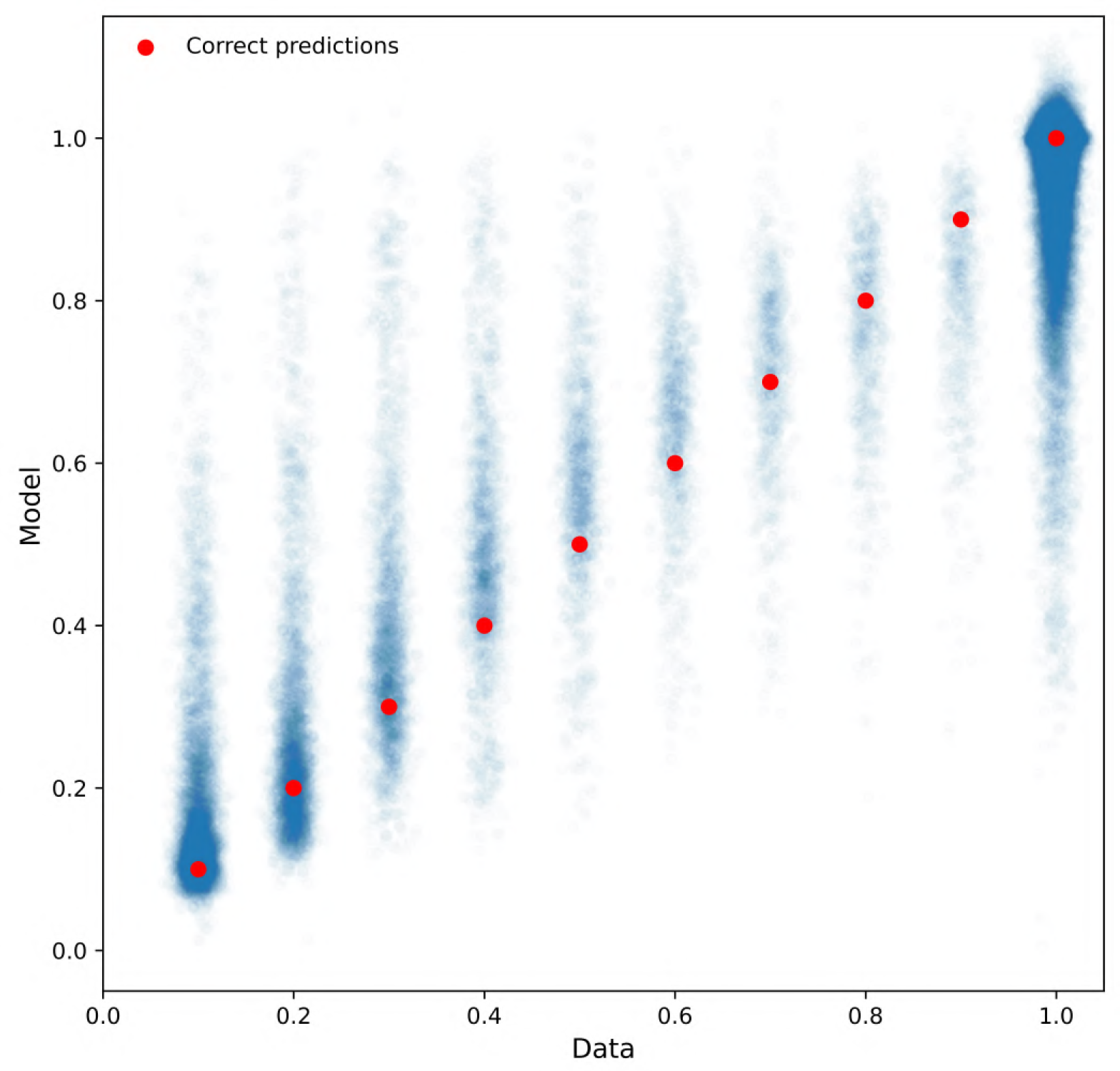
Gradient-boosted trees model performance on the test set – scatter plot. Here we plot the optimal dose from the data on the *x* axis against the model prediction on the *y* axis. We add some noise (in the *x*-direction) to the data, which should take values 0.1, 0.2,…, 1, so that it is clearer to see the density of points. The vast majority of data are at full dose (50,752 out of 70,000 in the test set), or at 0.1 or 0.2 (6560 + 3826 = 10386 out of 70,000). The red dots show what the correct predictions should be. The *R*^2^ value is 0.898.

## Appendix 3

### Parameter scan

#### Effect of the gamma distribution parameters

The effect of the gamma distribution mean and rate parameter on the resulting pathogen trait distribution.

We use the shape/rate parameterisation of the gamma distribution, so that if we have rate parameter *b* and shape parameter *a* = *μb* the PDF takes the following form:

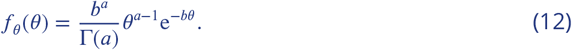

Also note that the trait mean can be expressed in terms of the gamma distribution mean *μ* and rate parameter *b* by:

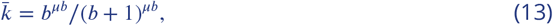

as derived below.

**Appendix 3 Figure 1.**
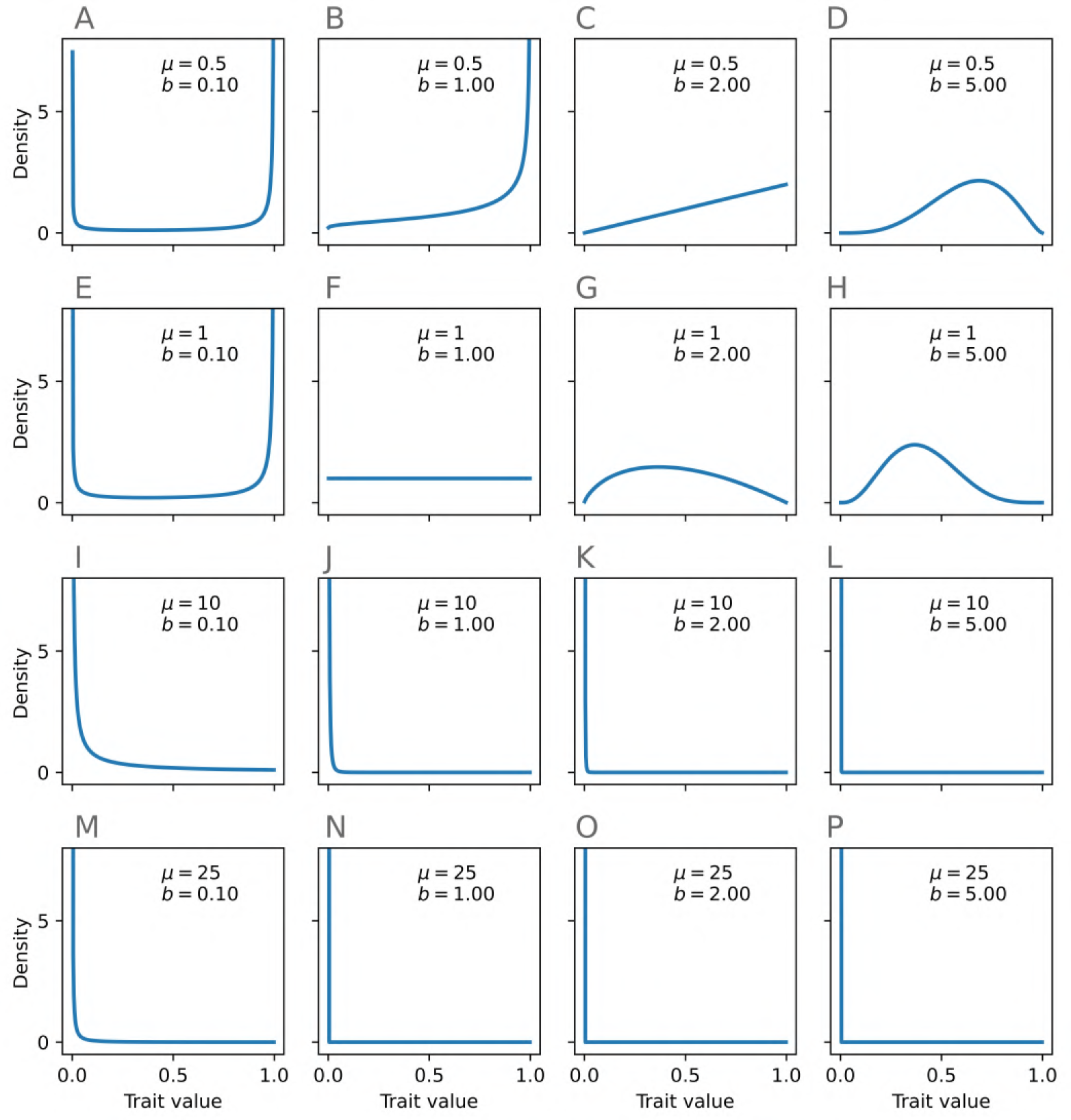
The effect of the gamma distribution parameters on the trait value distribution. Moving from left to right, the value of the gamma rate parameter *b* increases and the distribution variance decreases. For very low values of *b* this results in higher density at high *k* values as well as low *k* values. Moving from top to bottom, the value gamma distribution mean *μ* increases and the mean value of the trait value distribution decreases. To make the comparison clearer, we cut off the *y* axis at a value of 8 in all panels.

**Appendix 3 Figure 2.**
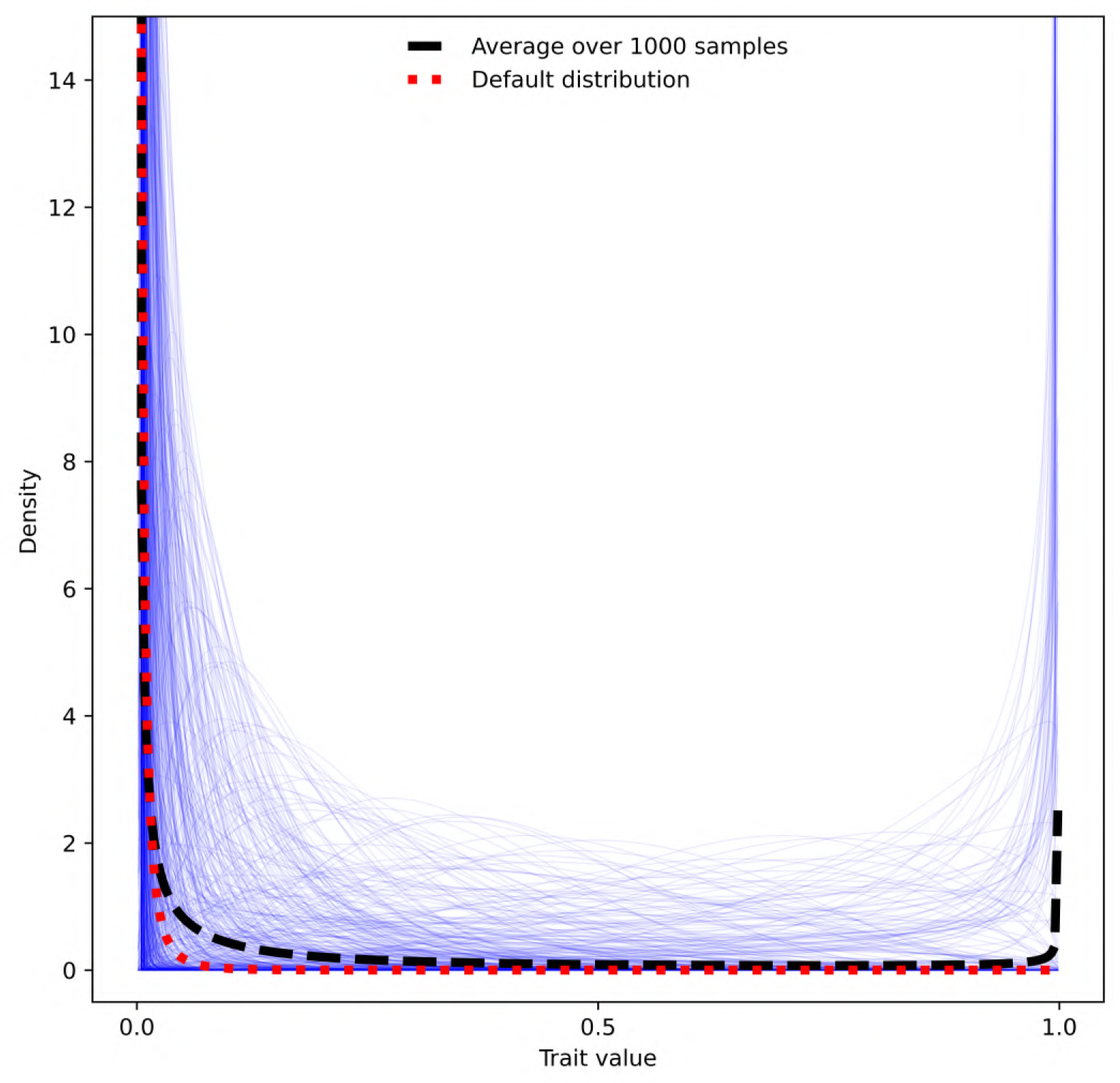
A visualisation of 1000 sampled distributions. Here we show 1000 distributions (thin blue curves) generated from values of the gamma distribution parameters *μ*, *b* sampled from the same distributions as in the parameter scan (Table 3. Although the density at *k* = 1 looks quite high when we take the mean across the 1000 distributions, this is skewed by a few extreme high values. For comparison we show the default trait distribution (as fitted to data in ***Taylor and Cunniffe*** (***2022a***)).

#### Derivation of the trait mean in terms of gamma distribution parameters

Suppose we have a gamma distribution for the curvature *θ* = −log(*k*), with mean *μ* and rate parameter *b*. Then *k* = e^−*θ*^. Denote the probability density function of *k* ∈ [0, 1] as *f*_*k*_ and of *θ* ∈ [0, ∞) as *f*_*θ*_. Then 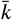 is defined as:

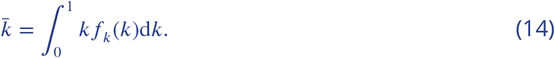

Note the standard result for transforming probability density functions:

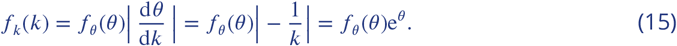

Also note that 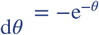. Then

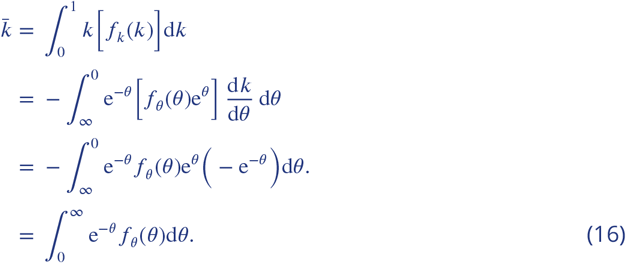

Then recall that the gamma distribution PDF with rate parameter *b* and shape parameter *a* = *μb* takes the following form:

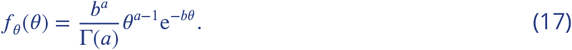

Then

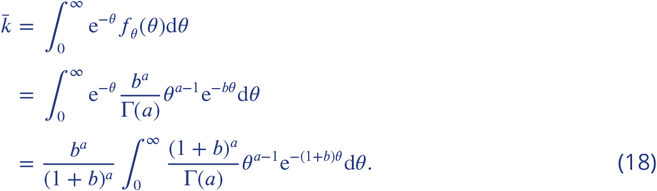

The final integral takes the form of a gamma distribution with shape parameter *a* and rate parameter *b* + 1, integrated from 0 to ∞. So this integral is equal to 1, and we find:

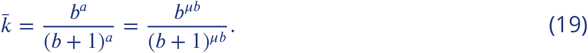

#### Example of distribution changes with different doses

For the model run from Figure 3A (main text), we show the pathogen distribution for different doses in years 1, 10, 20 and 30.

**Appendix 3 Figure 3.**
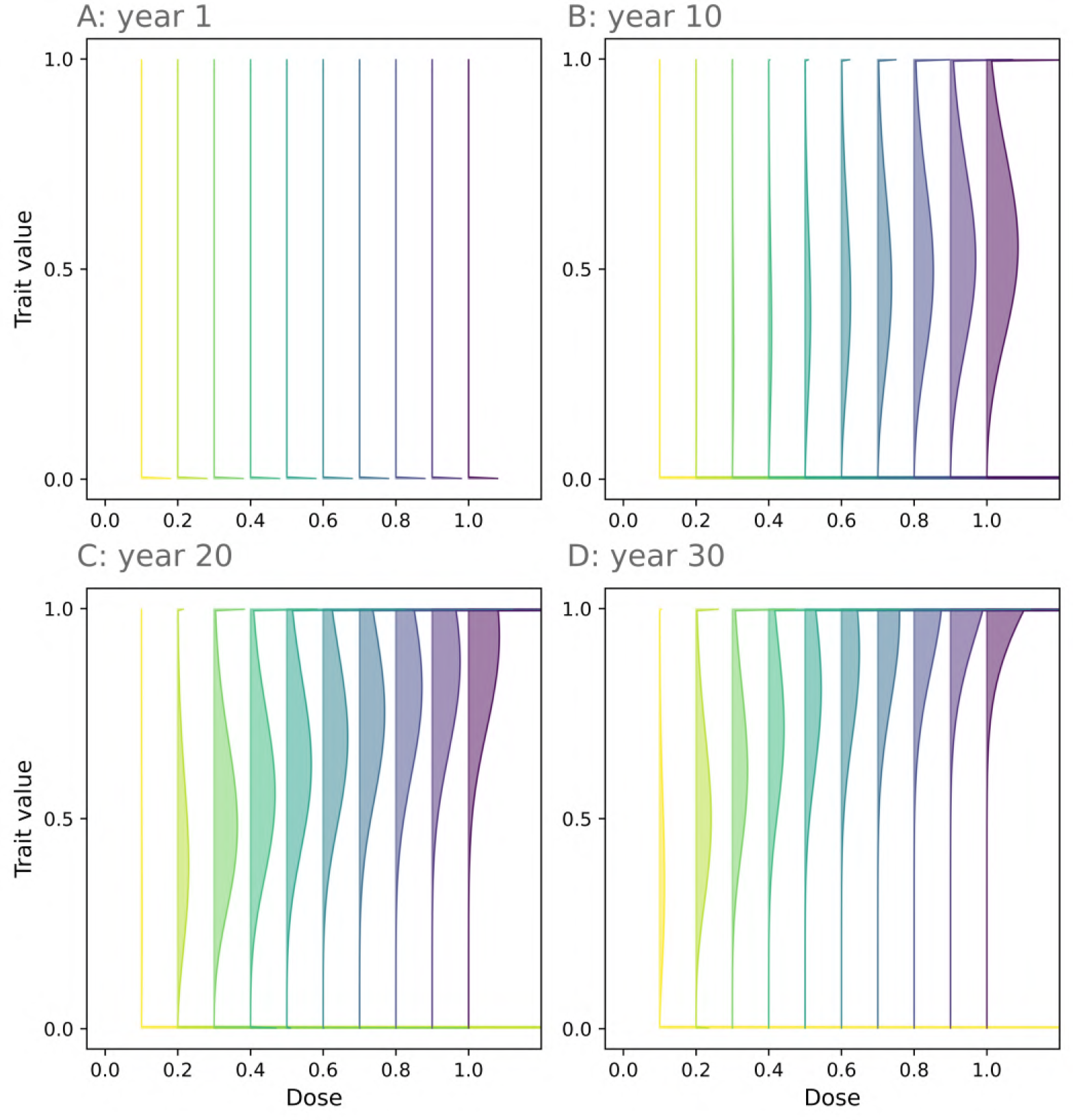
Pathogen distributions for different doses. Scaled pathogen distributions for the model run from Figure 3A (main text). Initially virtually all of the density is at *k* = 0 (**A**). As time goes on, the population becomes more resistant (**B**-**D**). This happens faster when the dose is higher. Parameter values are in Appendix 1 Table 2.

## Appendix 4

### Year 10 model

To explore the effect of the model parameters without the strong interaction with year, we fitted a model to data from year 10 only. The order of importance of the remaining model parameters stays the same, except for the mean effect at full dose *ν* and decay rate multiplier *M*_*d*_ features swapping order from 2nd and 3rd most important to 3rd and 2nd most important.

**Appendix 4 Figure 1.**
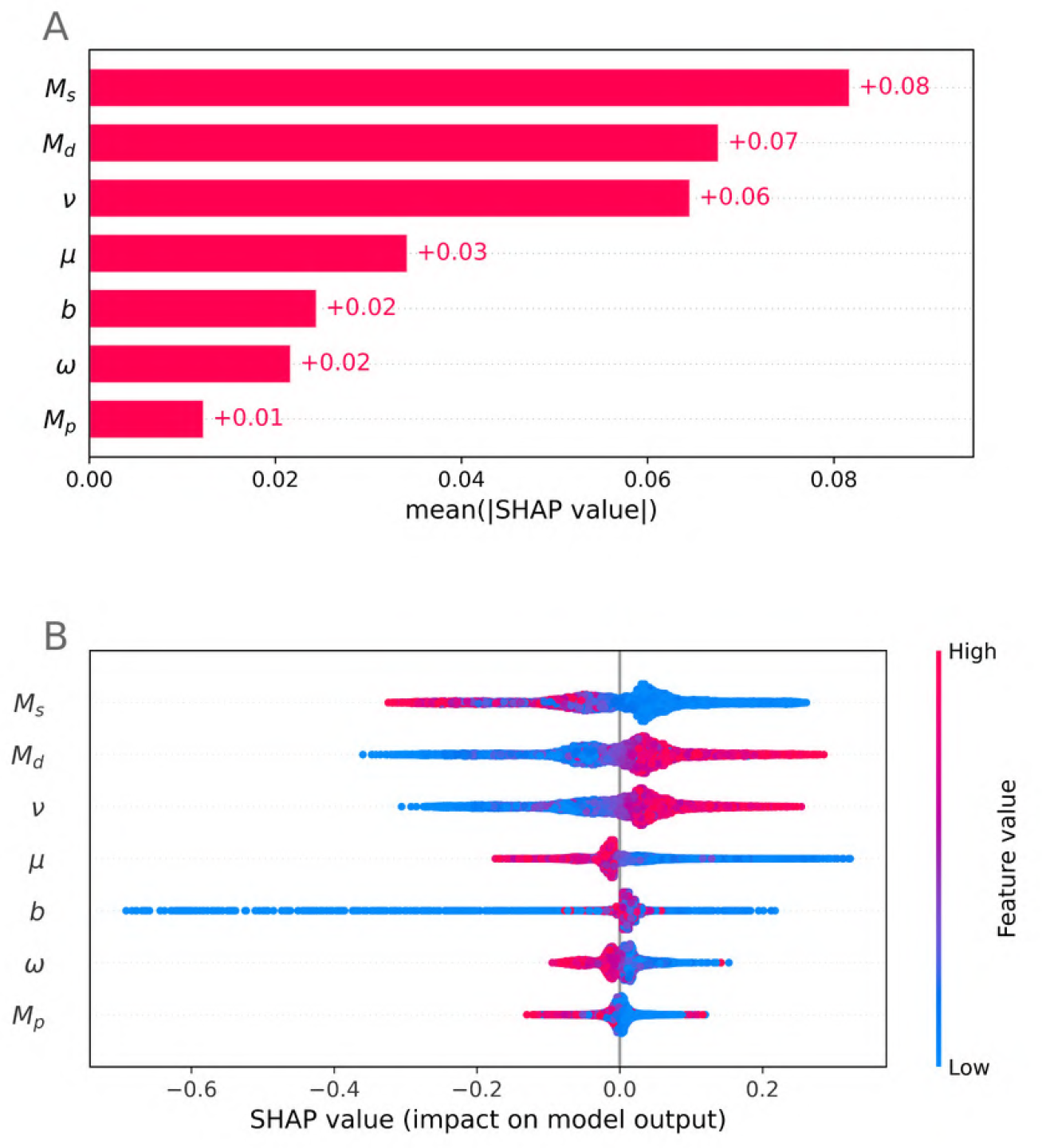
Year 10 model feature importance. The importance order remains the same for the features, except for *ν* and *M*_*d*_ which swap order. See Table 3 (main text) for a full explanation of model features and their ranges. Note that year is no longer a feature.

**Appendix 4 Figure 2.**
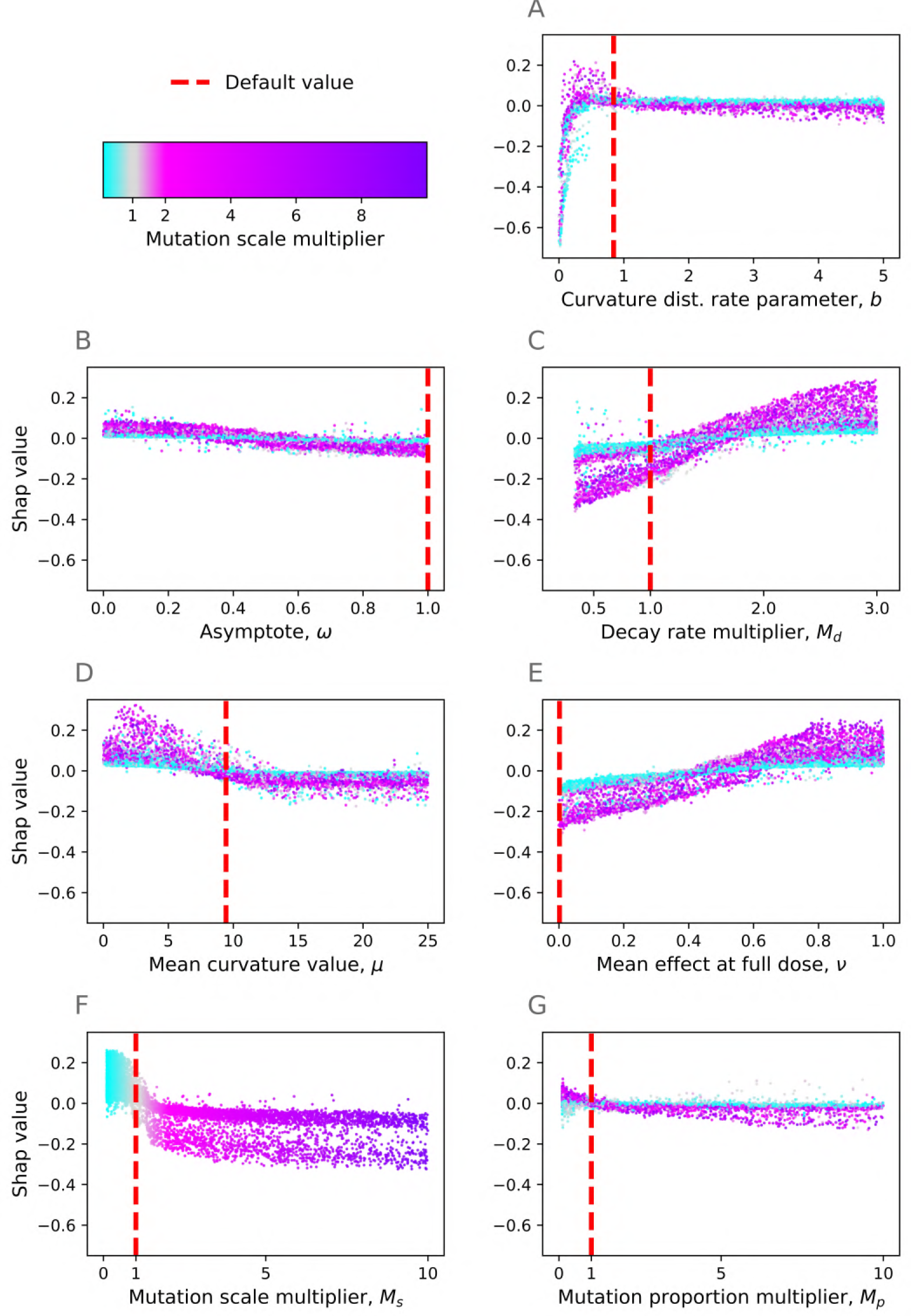
Year 10 model – effect of features. Here we plot the interaction with the mutation scale multiplier. Default values from ***Taylor and Cunniffe*** (***2022a***) are shown by the red dotted lines.

## Appendix 5

### Cumulative yield model

We wanted to explore which doses optimised the cumulative yield, i.e. for each year the dose which gave the highest total yield up until and including that year. The series of figures below replicate Figures 2, 3 and 5, 6 from the main text but focusing on the cumulative yield instead of the instantaneous yield in that year only. This involved fitting another gradient-boosted trees model and calculating the corresponding Shapley values. Since higher doses give higher yields initially, the optimal doses tend to be higher than when we considered the yield in each year separately/instantaneously in both the qualitative and quantitative resistance models (Figures 1, 2).

**Appendix 5 Figure 1.**
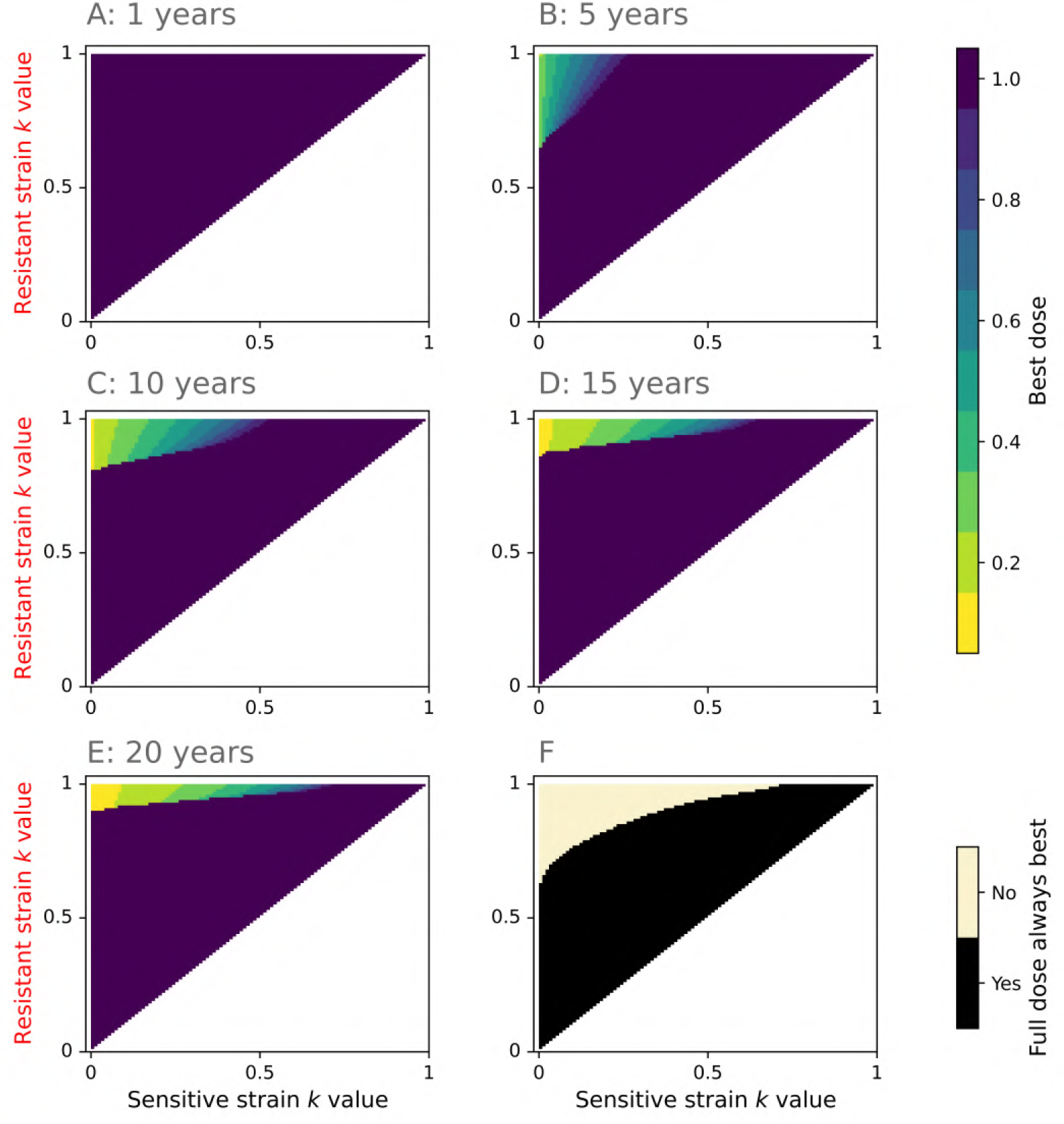
Optimal doses for cumulative yield in the qualitative resistance model. This figure is exactly analogous to Figure 2 (main text), but here the best dose is in terms of the cumulative rather than instantaneous dose. The region in which low doses are best is smaller (**F**), due to the initial benefit of using higher doses.

**Appendix 5 Figure 2.**
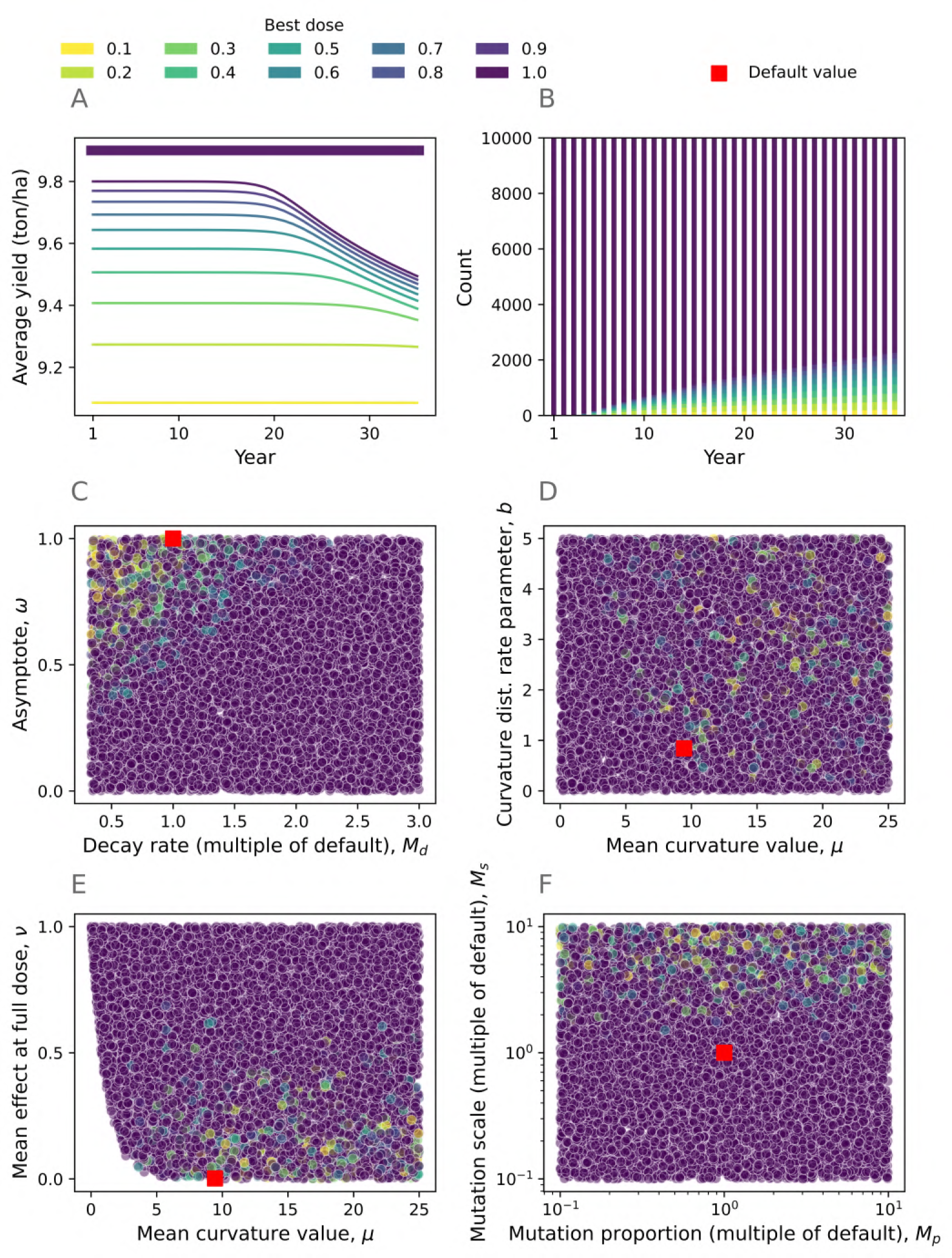
High doses are frequently best for cumulative yield in the quantitative resistance model. This figure is exactly analogous to Figure 3 (main text), but here the best dose is in terms of the cumulative rather than instantaneous dose. An example model run is shown in panel **A**. Here instead of the instantaneous yield we show the average yield, i.e. in each year the average yield obtained up until and including that year. As before, lower doses are often better for high asymptotes (*ω*) and low decay rates (low *M*_*d*_) (**C**). Again, The parameters in **D** have a less clear effect, although low doses are typically only better when *μ* takes higher values. Low mean effect at full dose *ν* often incentivises low doses (**E**). As before, the mutation scale plays a bigger role than the mutation proportion, with higher mutation scales often corresponding to lower doses being optimal (**F**). Parameter values for panel **A** are in Appendix 1 Table 2.

As we did in the main parameter scan, we fit a gradient-boosted trees model to the data and use Shapley values to try and interpret the effect of the model parameters on the dose which gave the optimal cumulative yield. We found that the most important features were very similar, with the 4 least important features taking exactly the same order. The 4 most important features are the same but in a different order, with year *Y* the fourth most important for cumulative yield (*M*_*s*_; *ν*; *M*_*d*_; *Y* in order of importance) rather than the second most important for instantaneous yield (*M*_*s*_; *Y*; *ν*; *M*_*d*_ in order of importance). Year is less important here most likely because full dose is best for cumulative yield in every year for many of the model runs in the parameter scan.

**Appendix 5 Figure 3.**
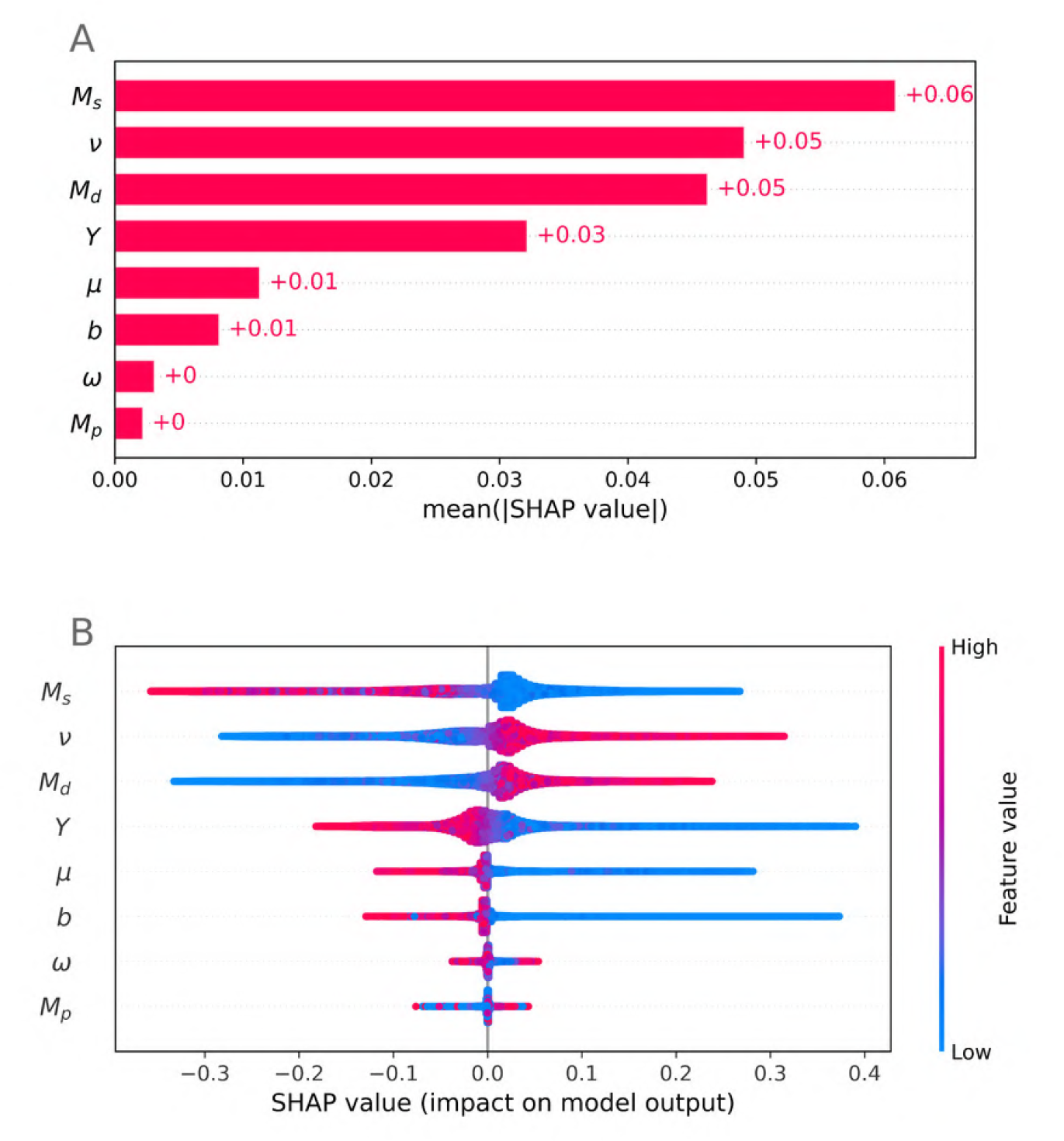
Which model features explain which dose optimises cumulative yield? The most important features are similar to those from Figure 5 (main text). Interestingly the decay rate multiplier *M*_*d*_ is more important than the year *Y* here - most likely because in many of the model runs full dose is the best in every year, so on average year has a relatively smaller effect than when optimising for instantaneously best yield.

**Appendix 5 Figure 4.**
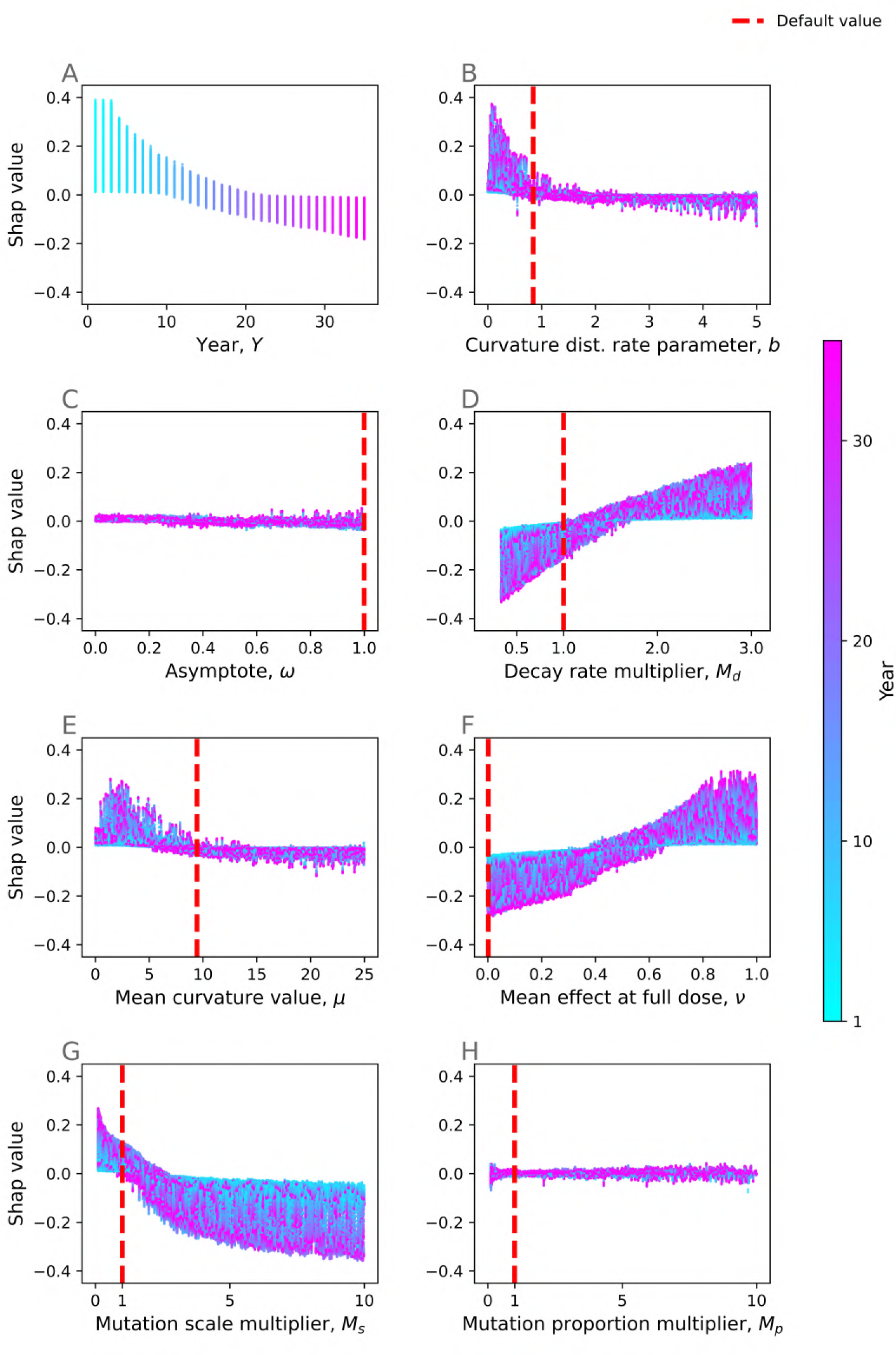
Which model features correlate with optimality of high doses? This figure is exactly analogous to Figure 6 (main text), but for cumulative yield rather than instantaneous yield. The parameters have a similar effect on the model output as in the instantaneous yield model.

## Appendix 6

### Partial resistance Type 1 model

To check whether the model results changed drastically if resistance were characterised in terms of the fungicide asymptote (partial resistance Type 1) rather that the fungicide curvature (partial resistance Type 2), we ran another parameter scan. This second scan is of interest since dose-convergence occurs for partial resistance Type 2, but not for partial resistance Type 1, and dose-convergence may influence whether selection for resistance is much stronger at high doses.

In this form of the model, differences in resistance in the population are characterised in terms of the fungicide asymptote *ω* rather than the curvature *θ*. This means that the distribution of *k* values relate to the asymptote (i.e. *k* = *ω*). The effect of the fungicide on the infection rate for a given strain with asymptote *k* and (constant) curvature *θ* is then:

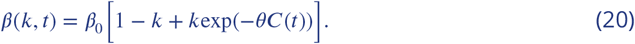

Note that the mean effect at full dose is then given by:

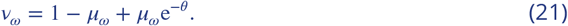

The initial distribution of asymptote (*ω*) values is given by a beta distribution since *ω* takes values in [0, 1] (Figure 1), whereas for the curvature (*θ*) we used a gamma distribution since the curvature takes values in [0, ∞). Recall the probability density function of a beta distribution with shape parameters *a*_*ω*_ and *b*_*ω*_:

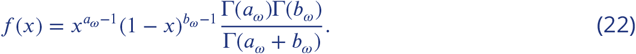

Also recall the standard result that the mean of a beta distribution is:

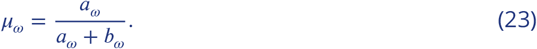

### Parameter ranges

The ranges for the parameter scan.

**Appendix 6 Table 1.** Parameters/features involved in the partial resistance Type 1 parameter scan. The first six are randomly and independently sampled for each parameter run, while the mean fungicide effect at full dose (*ν*_*ω*_) is calculated based on these, and year varies between 1 and 35 in each run. The first four parameters are sampled from a uniform distribution, and the two mutation multipliers are sampled from a log-uniform distribution. The multipliers for decay rate, mutation scale and mutation proportion relate to the default values for Λ, *σ*^2^ and *p*_*M*_ (Appendix 1).

| Symbol | Description | Range | Equation |
| --- | --- | --- | --- |
| $\mu_{\omega}$ | Pathogen initial distribution mean (asymptote) | [0.01, 1] | App. 6 Eqs. 22,23 |
| $b_{\omega}$ | Pathogen initial distribution shape parameter (asymptote) | [0.01, 30] | App. 6 Eq. 22 |
| $\theta$ | Fungicide curvature | [0.01, 30] | App. 6 Eq. 20 |
| $M_d$ | Fungicide decay rate multiplier | $\left[\frac{1}{3}, 3\right]$ | 4 (main text) |
| $M_s$ | Mutation scale multiplier | [0.1, 10] | 7 (main text) |
| $M_p$ | Mutation proportion multiplier | [0.1, 10] | 7 (main text) |
| $v_{\omega}$ | Mean fungicide effect at full dose | Depends on $\mu_{\omega}, \theta$ | App. 6 Eq. 21 |
| $Y$ | Year in model run | [1, 35] (but not sampled) | - |

### Effect of the beta distribution parameters

The effect of the beta distribution mean and rate parameter on the resulting pathogen trait distribution.

**Appendix 6 Figure 1.**
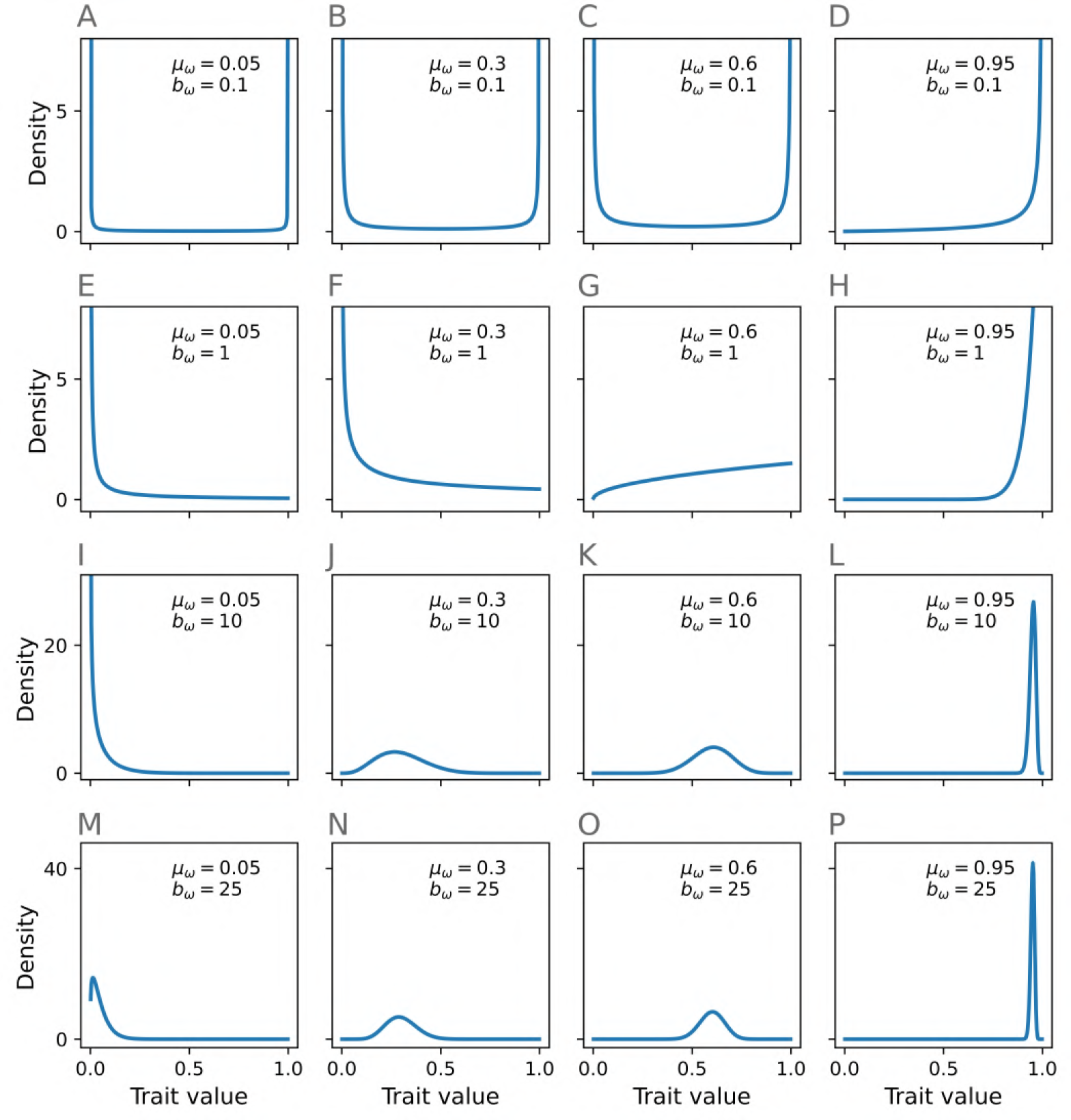
The effect of the beta distribution parameters on the trait value distribution in the partial resistance Type 1 scan. Moving from top to bottom, the value of the beta shape parameter *b*_*ω*_ increases. Moving from left to right, the value of the distribution mean *μ*_*ω*_ increases. To make the comparison clearer within each row, we cut off the *y* axis at a different value in each row. The beta distribution is extremely flexible which allows us to test out a wide range of possible initial distributions for the asymptote parameter *ω*.

### Results

Again, high doses were optimal frequently. However, lower doses were optimal more often than in the partial resistance Type 2 parameter scan (compare Figure 3, main text with Figure 2 below). Feature importances and variable impacts are shown in Figures 3, 4.

**Appendix 6 Figure 2.**
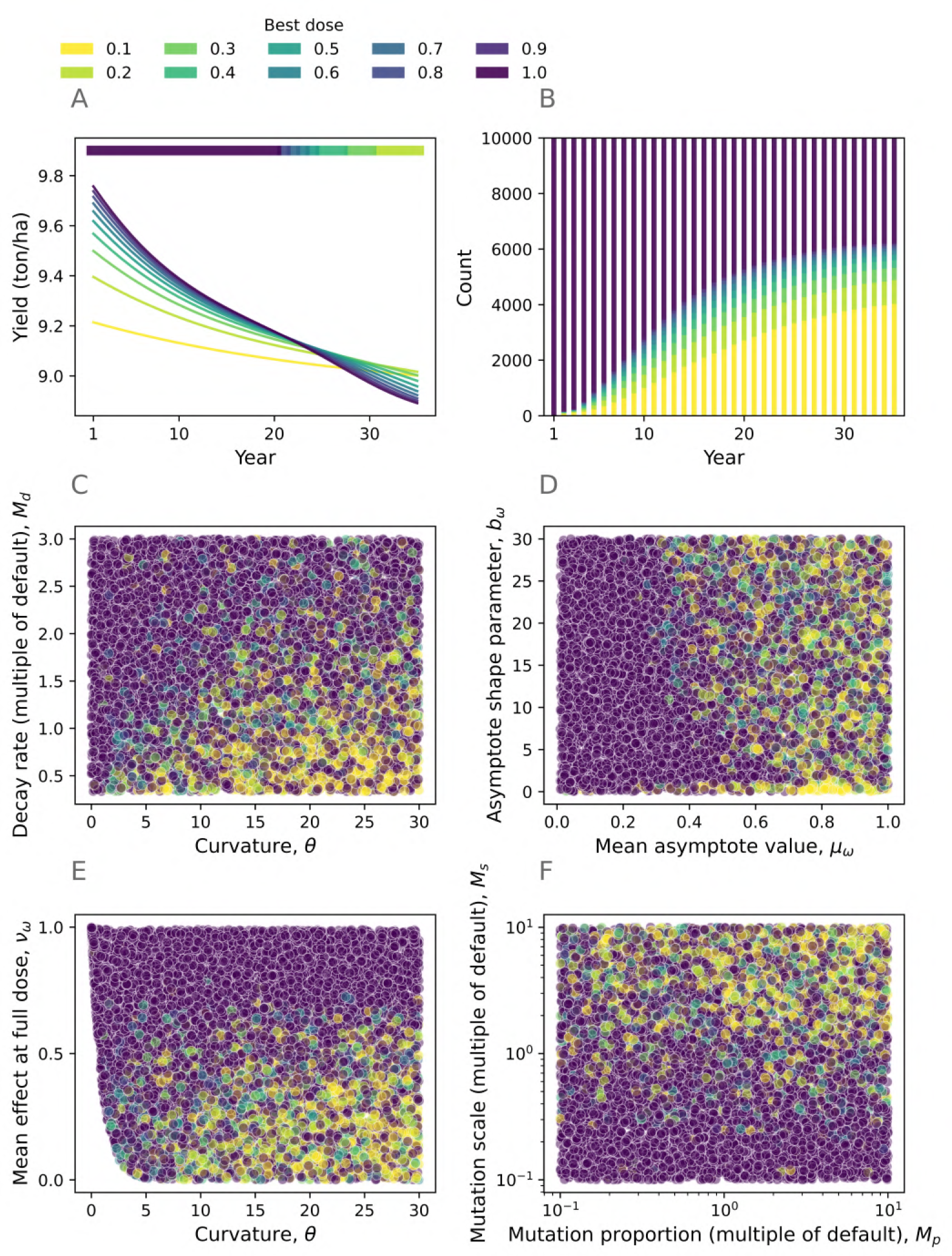
Partial resistance Type 1 parameter scan. This figure is analogous to Figure 3 (main text), but here applies to the partial resistance Type 1 parameter scan. Panel **A** shows one example model run from the ensemble of 10,000. The ‘best dose’ in any given year is the dose which gives the highest yield in that year, as shown by the colourbar at the top of panel **A**. Although full dose is best for over 50% of model runs in every year of the partial resistance Type 2 scan, in the partial resistance Type 1 scan by year 35 it is best in only 3808 of the 10,000 runs in the ensemble (**B**). Panels **C**-**F** relate to the 10th model year. High curvatures and low decay rates correlate with lower doses being best in year 10 (**C**). The asymptote shape parameter *b*_*ω*_ has little effect, but the higher mean asymptote values (*μ*_*ω*_) and mean effects at full dose *ν*_*ω*_ corresponding to stronger fungicides tend to correlate with lower doses being optimal (**D**,**E**). The mutation scale has a stronger effect than the mutation proportion (**F**). The results are similar to those in the other parameter scan (Figure 3, main text), but more exaggerated since low doses are better more frequently in this parameter scan. Parameter values in **A** (to 3 significant figures): *μ*_*ω*_=0.272, *b*_*ω*_=23.2, *θ*=17.1, *M*_*d*_ =1.55, *M*_*p*_=0.109, *M*_*s*_=1.72, *ν*_*ω*_=0.728.

**Appendix 6 Figure 3.**
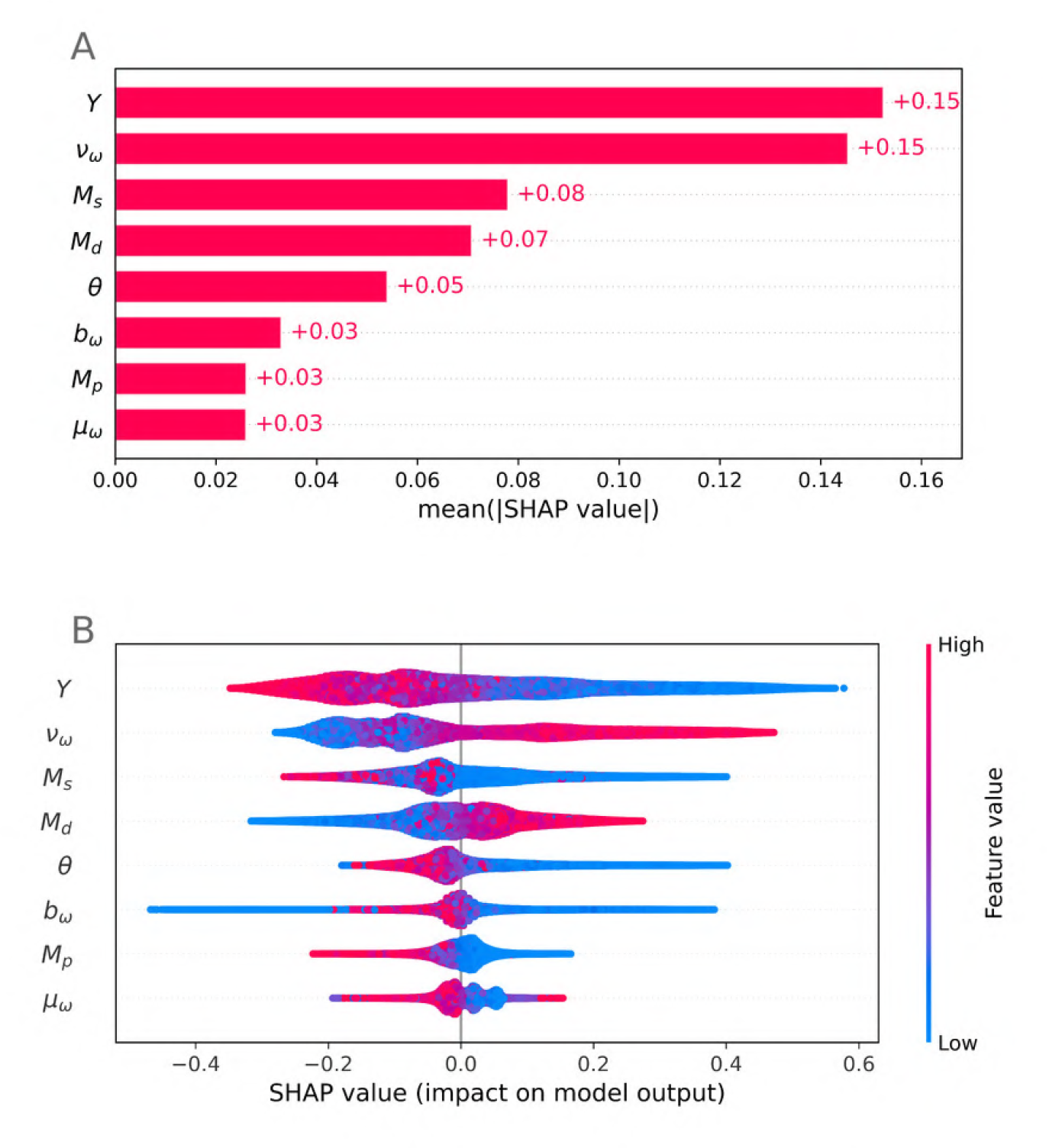
Which variables have the greatest impact on determining the optimal dose? This figure is analogous to Figure 5 (main text). The four most important features here are the year *Y*, the mean effect at full dose *ν*_*ω*_, the mutation scale multiplier *M*_*s*_ and the decay rate multiplier *M*_*d*_. These are the same four most important variables as in Figure 5 (main text), although the order is slightly different (in the main text it was in order of importance: *M*_*s*_; *Y*; *ν*; *M*_*d*_). The four other variables are of lower importance to the model output.

The optimal dose tends to decrease as year increases (**A**), and increase as the mean effect at full dose *ν*_*ω*_ increases (corresponding to reduced fungicide efficacy, **E**). The mean asymptote value *μ*_*ω*_ has a smaller effect, most likely because its impact on the model is captured via its effect on *ν*_*ω*_ (**B**). This matches the partial resistance Type 2 result (main text). The distribution shape parameter *b*_*ω*_ has a large but unpredictable effect when it is small (**C**). This is likely due to the wide range of distribution shapes that result from small values of *b*_*ω*_ (see Appendix 6 Figure 1). The decay rate variable interacts strongly with year *Y* – in later years rapid fungicide decay (large *M*_*d*_) correlates with optimality of lower doses, but in early years it correlates with optimality of higher doses (**D**). The opposite is true for slower decay rates (low *M*_*d*_). However, for partial resistance Type 2 (Figure 6, main text), increasing the decay rate tended to corrspond to increased optimal dose. Low curvature values *θ* correspond to increased optimal dose (**E**), similarly to the partial resistance Type 2 result (main text). As in the main text, mutation scale (**G**) has a bigger effect than mutation proportion (**H**). Mutation scale interacts strongly with year, having opposing effect in early years compared to later years (e.g. large mutation scales correspond to low doses being optimal in early years but high doses being optimal in later years, in contrast to the partial resistance Type 2 result - see Figure 6, main text). This is because if mutation scales are high, then by later years essentially all of the population is at very high *k* values, so virtually all control is lost by this point (Appendix 6 Figure 5). High doses can better control those remaining individuals away from *k* = 1, but by this point the control is minimal so the comparison becomes less important.

**Appendix 6 Figure 4.**
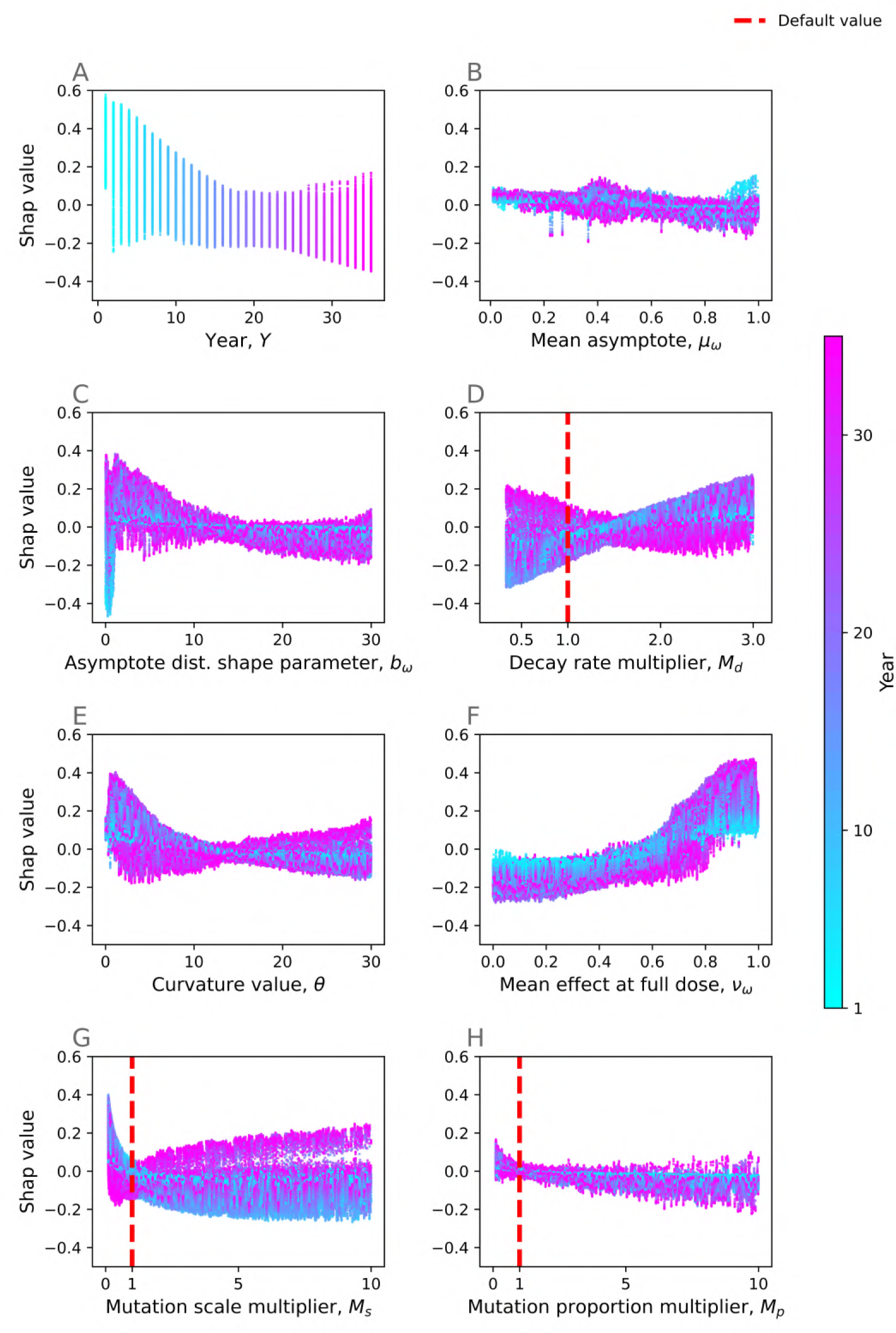
Which variables correlate with optimality of high doses? This figure is analogous to Figure 6 (main text). We show the effect of many of the model variables on the output. Each point corresponds to a single year from a single model run in the ensemble; *x*-values corresponds to a variable value, and *y*-values are the SHAP value for that feature. A high SHAP value corresponds to that variable changing the output to a the optimal dose being higher. The most important features (see Appendix 6 Figure 3) are the year (**A**), the mean effect at full dose (**E**), the mutation scale multiplier (**G**) and the decay rate multiplier (**H**). Some of the variables (e.g. mutation scale, mean effect and full dose, decay rate multiplier) interact strongly with the year – later years are shown in purple and early years in blue.

**Appendix 6 Figure 5.**
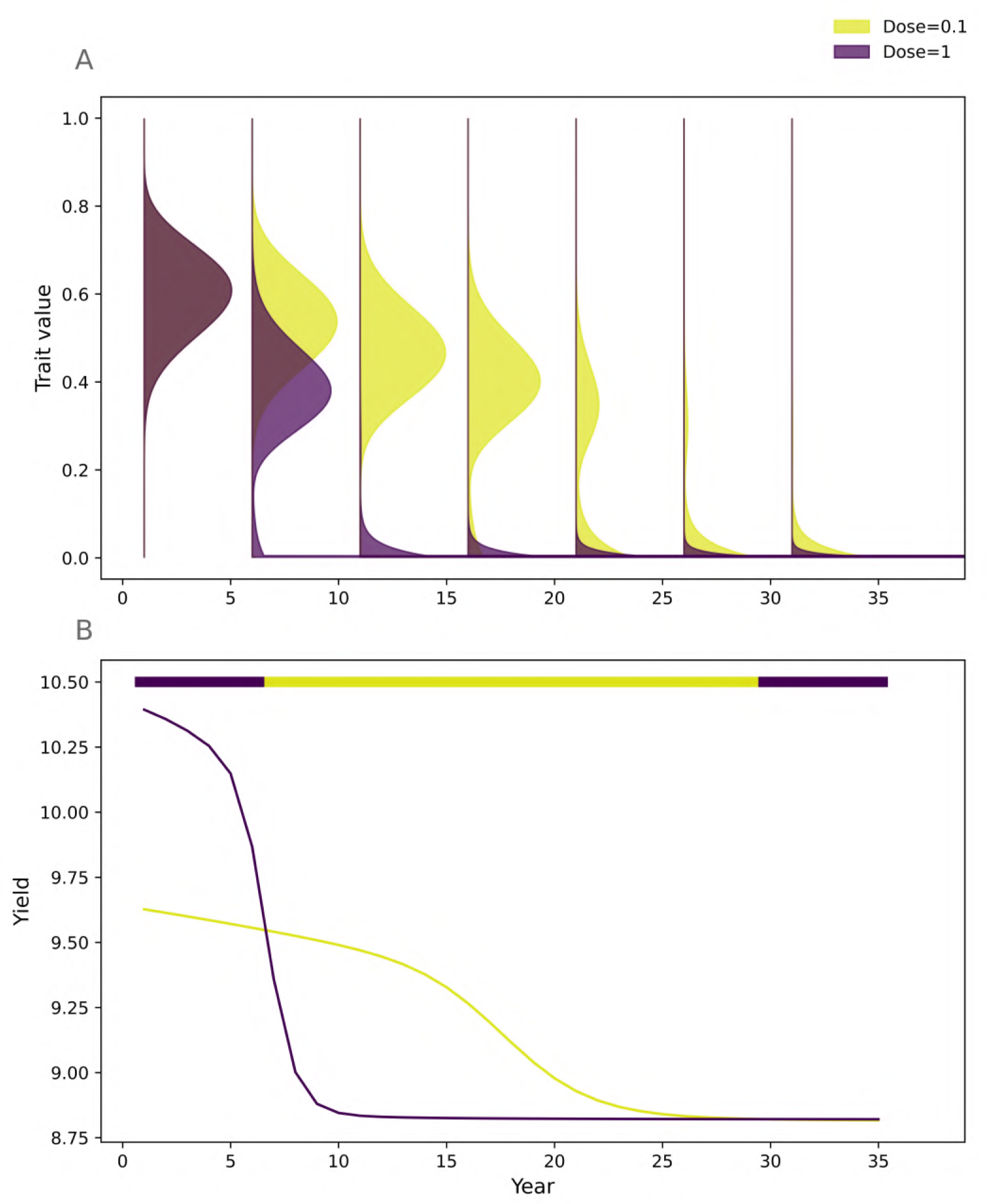
How could high mutation scales incentivise high doses? An example showing that if the mutation scale is high, resistant strains are rapidly selected for (**A**). The model suggests that high doses are best in later years (as shown by the colourbar showing the best dose, **B**). However by this point control for both doses is so ineffective that there is essentially no benefit to choosing either dose. Parameter values: *μ*_*ω*_=0.6, *b*_*ω*_=10.0, *θ*=10.0, *M*_*d*_ =1.0, *M*_*p*_=1.0, *M*_*s*_=10.0.

